# Maternal Circulating MiRNAs That Predict Infant FASD Outcomes Influence Placental Maturation

**DOI:** 10.1101/409854

**Authors:** Alexander M. Tseng, Amanda H. Mahnke, Alan B. Wells, Nihal A. Salem, Andrea M. Allan, Victoria H.J. Roberts, Natali Newman, Nicole A.R. Walter, Christopher D. Kroenke, Kathleen A. Grant, Lisa K. Akison, Karen M. Moritz, Christina D. Chambers, Rajesh C. Miranda, CIFASD

**Affiliations:** Department of Neuroscience and Experimental Therapeutics, Texas A&M University Health Science Center, Bryan, TX, USA; Clinical and Translational Research Institute, University of California San Diego, San Diego, CA, USA; Department of Pediatrics, University of California San Diego, San Diego, CA, USA; Department of Neurosciences, University of New Mexico, Albuquerque, NM, USA; Division of Reproductive and Developmental Sciences, Oregon National Primate Research Center, Oregon Health & Science University, Portland, OR, USA; Division of Neuroscience, Oregon National Primate Research Center, Oregon Health & Science University, Portland, OR, USA; Child Health Research Centre and School of Biomedical Sciences, the University of Queensland, 4072, Brisbane, Australia

**Keywords:** microRNA, placenta, EMT, trophoblast, fetal growth restriction

## Abstract

Prenatal Alcohol exposure (PAE), like other pregnancy complications, can result in placental insufficiency and fetal growth restriction, though the linking causal mechanisms are unclear. We previously identified 11 gestationally-elevated maternal circulating miRNAs that predicted infant growth deficits following PAE. Here, we investigated whether these _HEa_miRNAs contribute to the pathology of PAE, by inhibiting trophoblast epithelial-mesenchymal transition (EMT), a pathway critical for placental development. We now report for the first time, that PAE inhibits expression of placental pro-EMT pathway members in both rodents and primates, and that _HEa_miRNAs collectively, but not individually, mediate placental EMT inhibition. _HEa_miRNAs collectively, but not individually, also inhibited cell proliferation and the EMT pathway in cultured trophoblasts, while inducing cell stress, and following trophoblast syncytialization, aberrant endocrine maturation. Moreover, a single intra-vascular administration of the pooled murine-expressed _HEa_miRNAs, to pregnant mice, decreased placental and fetal growth and inhibited expression of pro-EMT transcripts in placenta. Our data suggests that _HEa_miRNAs collectively interfere with placental development, contributing to the pathology of PAE, and perhaps also, to other causes of fetal growth restriction.

**Summary:** Maternal gestational circulating microRNAs, predictive of adverse infant outcomes including growth deficits, following prenatal alcohol exposure, contribute to placental pathology by impairing the EMT pathway in trophoblasts.

## Introduction

Prenatal alcohol exposure (PAE) is common (1–3). Between 1.1-5% of school children in the United States are conservatively estimated to have a Fetal Alcohol Spectrum Disorder (FASD, (4)). Consequently, FASD, due to PAE, is the single largest cause of developmental disabilities in the US and worldwide (5), and a co-morbid factor in a number of other prevalent developmental neurobehavioral disabilities including attention deficit/hyperactivity and autism spectrum disorders (6).

PAE can result in decreased body weight, height and/or head circumference in infants. Consequently, infant growth deficits are a cardinal diagnostic feature for Fetal Alcohol Syndrome (FAS, (7)), which represents the severe end of the FASD continuum. However, though well-recognized as a diagnostic feature, the mechanistic linkage between PAE and growth restriction remains unclear. In 2016, as part of our effort to identify maternal diagnostic biomarkers of the effect of PAE, we reported that elevated levels of 11 distinct microRNAs (miRNAs) in maternal circulation during the 2^nd^ and 3^rd^ trimesters distinguished infants who were affected by *in-utero* alcohol exposure (Heavily Exposed Affected: HEa) from those who were apparently unaffected at birth by PAE (Heavily Exposed Unaffected: HEua), or those who were unexposed (UE) (8). In that study, we predicted, based on bioinformatics analyses, that these _HEa_miRNAs (MIMAT0004569 [hsa-miR-222-5p], MIMAT0004561 [hsa-miR-187-5p], MIMAT0000687 [hsa-mir-299-3p], MIMAT0004765 [hsa-miR-491-3p], MIMAT0004948 [hsa-miR-885-3p], MIMAT0002842 [hsa-miR-518f-3p], MIMAT0004957 [hsa-miR-760], MIMAT0003880 [hsa-miR-671-5p], MIMAT0001541 [hsa-miR-449a], MIMAT0000265 [hsa-miR-204-5p], MIMAT0002869 [has-miR-519a-3p]), could influence signaling pathways crucial for early development, particularly the epithelial-mesenchymal transition (EMT) pathway.

Placental development involves maturation of cytotrophoblasts at the tips of anchoring villi into invasive extravillous trophoblasts, as well as fusion of cytotrophoblasts into multinucleate, hormone-producing syncytiotrophoblasts (9). Maturation into extravillous trophoblasts, which invade the maternal decidua and remodel the uterine spiral arteries into low-resistance high-flow vessels that enable optimal perfusion for nutrient and waste exchange, requires cytotrophoblasts to undergo EMT (10). Impaired placental EMT, as well as orchestration of the opposing mesenchymal-epithelial transition pathway, has been found in conditions resulting from placental malfunction, primarily preeclampsia (11–16). While there have been no previous studies directly investigating the effects of PAE on placental EMT, a rodent study demonstrated that PAE, during a broad developmental window, reduced the number of invasive trophoblasts within the mesometrial triangle, a region of the uterine horn directly underlying the decidua (17). Furthermore, both human and rodent studies have found PAE disrupts placental morphology, and interferes with cytotrophoblast maturation, as with preeclampsia (18–21). Disrupted trophoblast maturation, seen in these conditions, is associated with aberrant expression of placental hormones, primarily human chorionic gonadotropin (hCG) (22–25).

Our study is the first to report that PAE interferes with expression of core placental EMT pathway members. Using rodent and primate models of gestation, as well as complementary miRNA overexpression and knockdown studies *in vitro*, we also provide evidence that _HEa_miRNAs, which predict infant growth deficits due to PAE, collectively but not individually, mediate PAE’s effects on placental EMT through their effects on cytotrophoblast maturation and stress. In a mouse model of pregnancy, a single combined exposure to the murine-expressed _HEa_miRNAs, resulted in placental EMT inhibition, and diminished placental and fetal growth. Collectively, these data suggest that elevated _HEa_miRNAs may represent an emergent maternal stress response that triggers fetal growth restriction, though sub-groups of _HEa_miRNAs may compete to protect against the loss of EMT. Moreover, most members of the group of_HEa_miRNAs, have also been implicated in other placental insufficiency and growth restriction syndromes, giving rise to the possibility that growth restriction syndromes may share common etiological mediators.

## Results

### _HEa_miRNAs are implicated in placental-associated pathologies

Given our prediction that _HEa_miRNAs interfere with signaling pathways governing fetal and placental development (8), we conducted a literature review of reports on _HEa_miRNA levels in gestational pathologies caused by poor placentation (26–28). Surprisingly, placental and plasma levels of 8 out of 11 _HEa_miRNAs were significantly dysregulated in one or more of these gestational pathologies with expression of the majority of these 8 miRNAs altered in both fetal growth restriction and preeclampsia (Figure 1A) (29–49), both of which are characterized by poor placental invasion (50–56).

**Figure 1:**
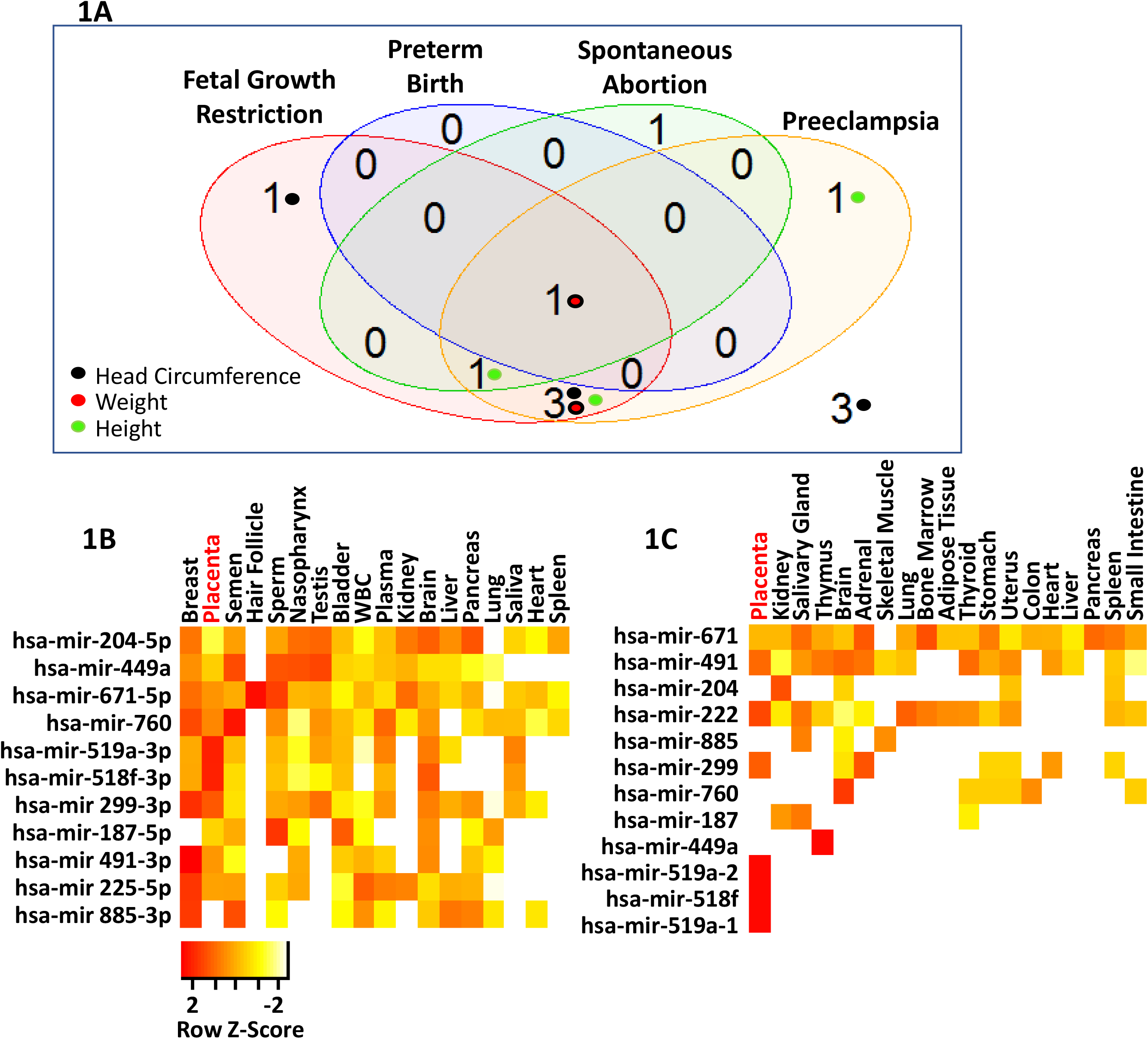
_HEa_miRNAs are placentally enriched and associated with gestational pathologies. **A)** Venn diagram on number of _HEa_miRNAs reported to be associated with different gestational pathologies. Inset colored circles represent the corresponding sex and gestational age-adjusted growth parameters these miRNAs were correlated with. Of the 22 studies queried, 11 (50%) utilized unbiased screenings for miRNA expression. **B)** Heatmap of mature _HEa_miRNA expression and C) pri-_HEa_miRNA expression across different tissues resulting from secondary analysis of publicly available RNA-sequencing data. Legend depicts row-centered Z-score.

### _HEa_miRNAs explain variance in infant growth outcomes due to PAE

Given the association of individual _HEa_miRNAs with gestational pathologies, we sought to determine if circulating _HEa_miRNAs levels could explain the variance in sex and gestational age-adjusted neonatal height, weight and head circumference in our Ukrainian birth cohort, which are growth measures sensitive to in utero environment (57). We found that 8 of the _HEa_miRNAs, each significantly explained between 7 to 19% of infant variation in these growth measures (Table 1). Furthermore, 7 of these miRNAs were also associated with fetal growth restriction and preeclampsia as identified by our literature review (Figure 1A). Interestingly, a multivariate statistical regression model that accounted for levels of all 11 _HEa_miRNAs together, explained a far greater proportion of infant variance, between 24-31%, in all three growth measures than accounting for them individually (Supplementary Table 2) suggesting _HEa_miRNAs collectively account for the variance in infant growth outcomes.

### _HEa_miRNAs are transcribed preferentially in the placenta

Data extracted from publicly available gene expression profiling datasets (58) show that _HEa_miRNAs as well as their unprocessed precursor transcripts, _HEa_pri-miRNAs, are enriched in placenta compared to other tissues, suggesting that the placenta itself transcribes these miRNAs and may be a significant contributory tissue to maternal circulating _HEa_miRNAs (Figures 1B and C). Moreover, since _HEa_miRNAs are also associated with gestational pathologies caused by poor placental invasion, these _HEa_miRNAs may also contribute to the placental response to PAE. We therefore assessed in rodent and primate models, whether PAE could result in impaired EMT, and if _HEa_miRNAs could explain the effects of PAE on placental EMT-associated gene expression.

### _HEa_miRNAs moderate placental EMT impairment in PAE models

EMT, in trophoblasts, is characterized by the disappearance of epithelial markers like E-Cadherin and the appearance of the mesenchymal markers like the intermediate filament, Vimentin, a process that is controlled by the expression of key mesenchymal determination transcription factors, Snail1 and 2 and TWIST, as extensively described (10, 14, 15, 59–62). These five markers have been used extensively to assess EMT in a variety of model systems, so our studies utilized these markers to assess the effects of alcohol and _HEa_miRNAs on trophoblast EMT.

In the first analysis, using a murine model of PAE that mimicked moderate to binge-type alcohol consumption throughout early and mid-pregnancy, we fractionated GD14 placenta into three zones: the cytotrophoblast and syncytiotrophoblast rich labyrinth zone, the glycogen and spongiotrophoblast rich junctional zone, and the decidual zone comprising the endometrial contribution to the placenta (Figure 2A). Multivariate analysis of variance (MANOVA) for expression of these five core genes in the EMT pathway within placental trophoblasts, revealed a significant effect of ethanol exposure on EMT pathway member expression selectively within the labyrinth zone (Pillai’s trace statistic, F_(5,21)_=6.85, p<0.001, Figure 2B) but not within the junctional or decidual zones. Post-hoc univariate ANOVA indicated ethanol exposure specifically elevated *CDH1* (F_(1,25)_=7.452, p=0.011), which encodes epithelial E-Cadherin, whereas expression of the pro-mesenchymal transcription factor *SNAI1* was significantly reduced (F_(1,25)_=21.022, p=0.0001). We also observed a significant interaction between fetal sex and PAE on expression of *SNAI2* (F_(1,25)_=2.18, p=0.047) and a trend towards decreased expression of the terminal mesenchymal marker *VIM* (Vimentin, F_(1,25)_=2.749, p=0.11), while there was no effect on *TWIST* expression (Figures 3A-3E). Consistent with our gene expression data, E-Cadherin protein levels were significantly elevated in the labyrinth zone of PAE placenta (F_(1,24)_=31.63, p=0.0005), while not in the junctional or decidual zones (Figure 3F and Supplementary Figures 3A and B). However, when we controlled for expression of the 8 mouse homologues of _HEa_miRNAs as a covariate, using multivariate analysis of covariance (MANCOVA), ethanol’s effect on EMT became marginally nonsignificant (Pillai’s trace, F_(5,21)_=2.713, p=0.068) (Figure 2C), suggesting that these miRNAs partially mediate effects of PAE on EMT pathway members in mice. Interestingly, PAE limited to the peri-conceptional period in rats also influenced expression of EMT core transcripts (Supplementary Figures 2B and 4A-4E).

**Figure 2:**
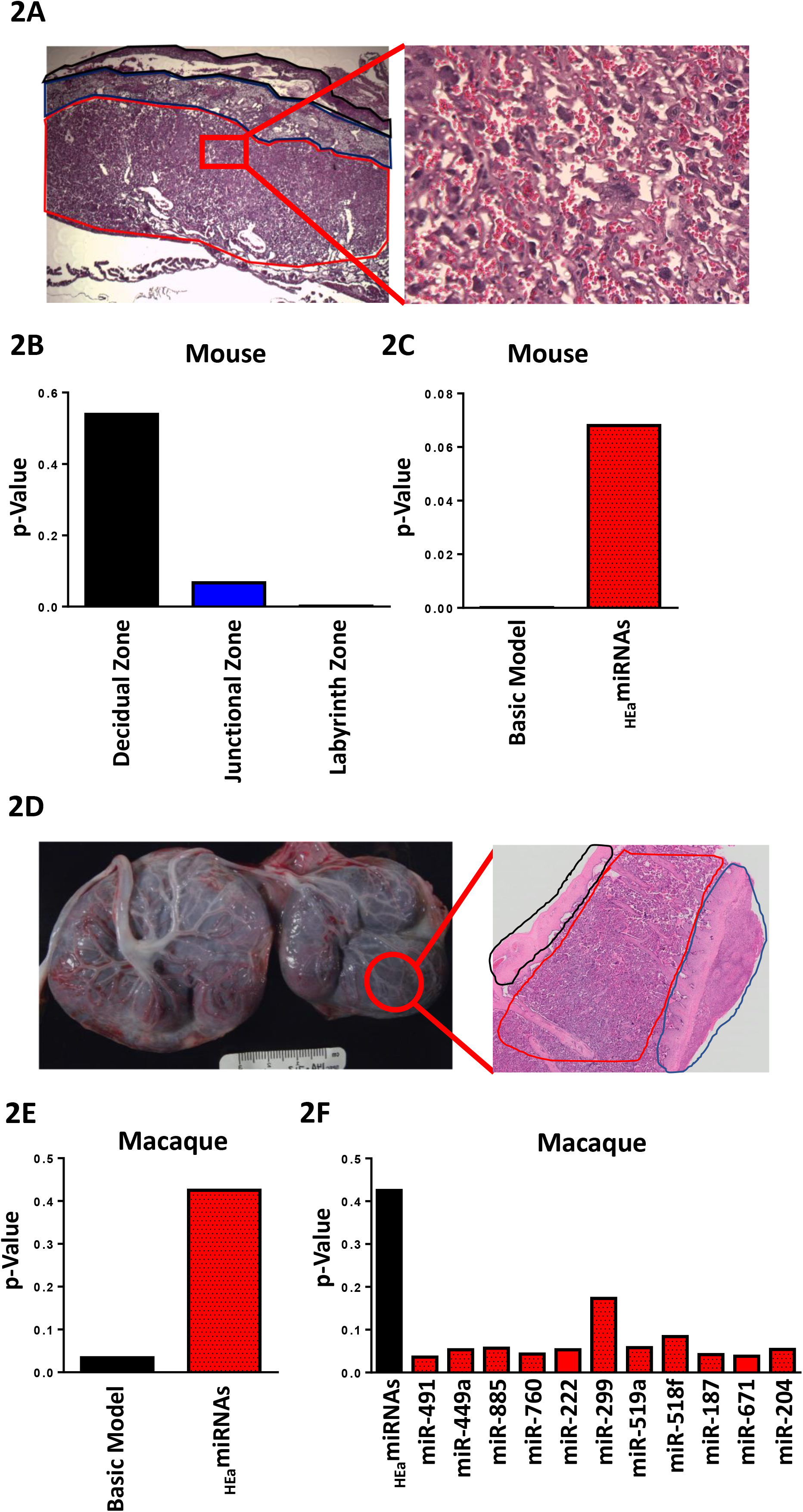
_HEa_miRNAs mediate the effect of PAE on EMT pathway members in mouse and macaque placentas. **A)** Histological image of GD14 mouse placenta. Outlined in red is the labyrinth zone, blue is the junctional zone, black is the decidual zone. Inset is a high magnification image of the labyrinth zone. **B)** MANOVA of gene expression of core EMT pathway members in different regions of the mouse placenta in control and PAE mice (n=29 samples). **C)** MANCOVA of gene expression of core EMT pathway members in the mouse placental labyrinth zone before (Basic Model) and after accounting for the expression of _HEa_miRNAs (n=29 samples). **D)** Gross anatomy photograph of the primary (left) and secondary (right) lobes of a GD135 macaque placenta. Outlined in red is an individual cotyledon from the secondary lobe. Inset is a full thickness hematoxylin and eosin stained histological section of a representative cotyledon with the fetal membranes outlined in black, villous tissue outlined in red and maternal decidua in blue. **E)** MANCOVA of gene expression of core EMT pathway members in placental cotyledons of PAE and control macaques, accounting for the expression of _HEa_miRNAs collectively (n=23 samples). **F)** MANCOVA of gene expression of core EMT pathway members in macaque placentas after accounting for expression of _HEa_miRNAs individually (n=23 samples).

**Figure 3:**
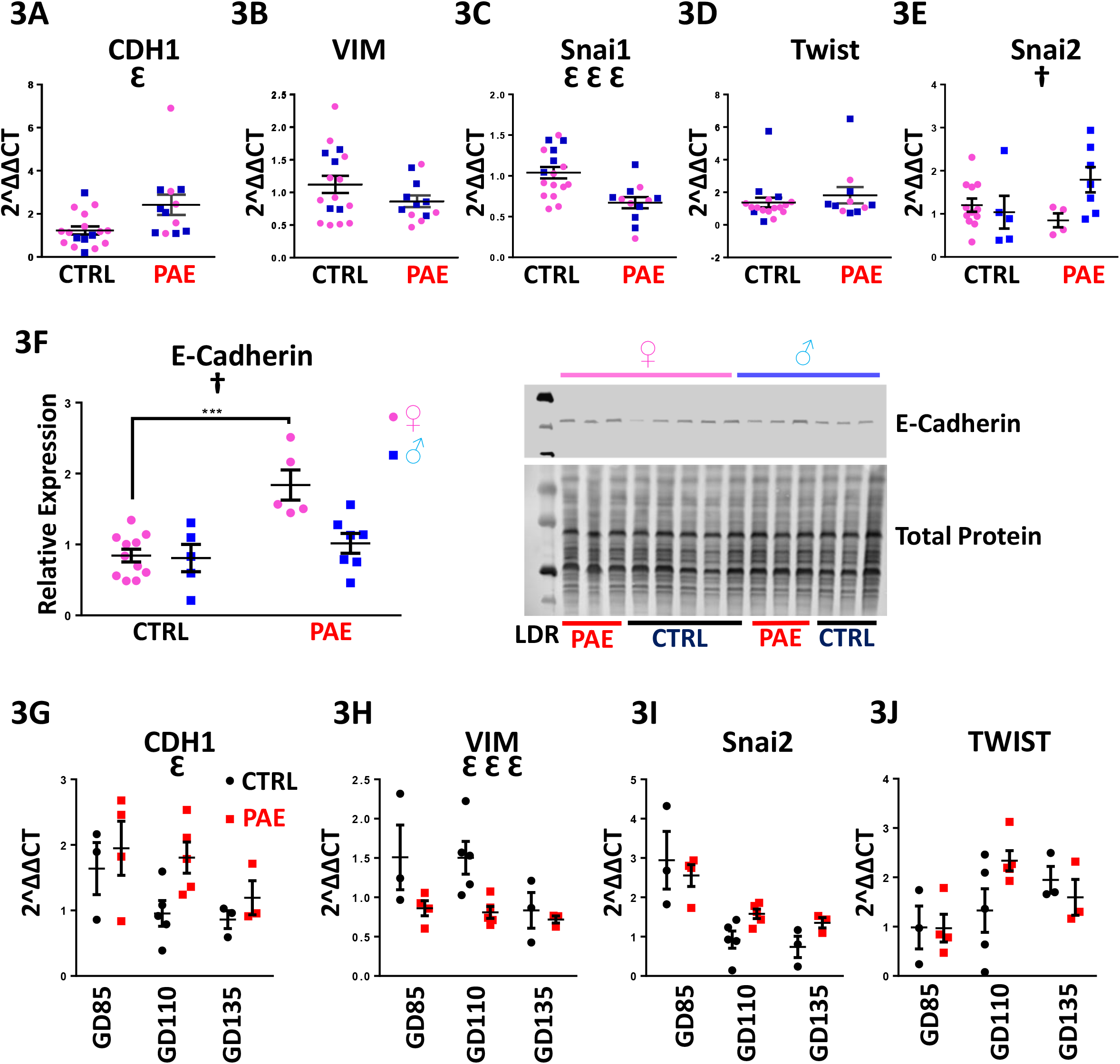
PAE interferes with the EMT pathway in mouse and macaque placentas. Expression of **A)** *CDH1*, **B)** *VIM*, **C)** *SNAI1*, **D)** *TWIST*, and **E)** *SNAI2* in the placental labyrinth zone of PAE and control mice (n=5-12 samples per group). **F)** Densitometric quantification of E-Cadherin expression in the labyrinth zone of PAE and control mice as well as representative blot of E-Cadherin expression and total protein expression (right, n=5-12 samples per group). Expression of **G)** *CDH1*, **H)** *VIM*, **I)** *SNAI2*, and **J)** *TWIST* transcripts in PAE and Control macaque placental cotyledons (n=3-5 samples per group). Results are expressed as the mean ± SEM, LDR=Molecular Weight Ladder; ANOVA: significant main effect of PAE ⍰, significant interaction effect (sex by PAE, [^†^p<0.05]). For post-hoc analysis, ***p<0.001 by Tukey’s HSD.

To determine if PAE’s effects on EMT pathway members in placenta are broadly conserved throughout mammalian evolution, we adopted a non-human primate (macaque) model of moderate to binge-type alcohol consumption. Placental tissues were isolated from GD85, GD110, and GD 135 placenta (Figure 2D), which spans the human equivalent of mid-second to mid-third trimester (Supplementary Figure 2C). There was a significant effect of ethanol exposure on expression of core EMT mRNA transcripts by MANOVA (Pillai’s trace statistic, F_(4,9)_=4.229 p=0.045, Figure 3B). Consistent with our findings in mouse, post-hoc univariate ANOVA indicated that in primate placenta, ethanol exposure significantly increased *CDH1* expression (F_(1,12)_=4.866, p=0.048) whereas *VIM* expression was significantly reduced (F_(1,12)_=12.782, p=0.0004), suggesting that, as in the mouse, PAE also impairs EMT in the primate placenta. Interestingly, there was no effect on *SNAI2* or *TWIST* expression (Figures 3G-3J). As in mice, accounting for expression of _HEa_miRNAs together as a covariate abolished the significant effect of PAE on EMT, though to a greater degree than mice (Pillai’s trace, F_(1,1)_=1.605, p=0.425, Figure 2E). Interestingly, accounting for expression of individual _HEa_miRNAs did not explain the effects of PAE on placental EMT, suggesting that _HEa_miRNAs act in concert to mediate the effect of PAE on EMT in primate placenta (Figure 2F).

Collectively, our data suggests PAE induced impairment of EMT in the trophoblastic compartment of placentae is conserved between rodents and non-human primates and that_HEa_miRNAs, particularly in primates, may moderate the effect of PAE on placental EMT.Consequently, subsequent studies focused on the collective role of _HEa_miRNAs, either on basal or on alcohol-influenced placental trophoblast growth, invasion, and the maturation of physiological function.

### _HEa_miRNAs impair EMT in a model of human cytotrophoblasts

To investigate whether _HEa_miRNAs collectively interfere with the EMT pathway, as suggested by our *in vivo* data, we examined the effects of transfecting _HEa_miRNA mimics and antagomirs into BeWO cytotrophoblasts (Figure 4A). We initially overexpressed each of the 11 _HEa_miRNAs individually, to determine whether any of them could influence the EMT pathway. We did not observe any significant effects (Supplementary Figure 5), consistent with our findings in the primate PAE model that individual miRNAs did not explain the effects of ethanol on EMT. In contrast, transfection of pooled _HEa_miRNAs into cytotrophoblasts significantly increased *CDH1* expression (F_(1,36)_=30.08, p<0.0001). Interestingly, expression of the pro-mesenchymal transcription factors *TWIST* and *SNAI1* were also significantly reduced, but only in the context of concomitant 320 mg/dL ethanol treatment, pointing to an interaction effect between _HEa_miRNAs and ethanol (F_(1,36)_= 5.650 and 5.146 respectively, p=0.023 and p=0.029, Figures 4B-E). Consistent with our qPCR data, transfection of _HEa_miRNAs also significantly increased E-cadherin protein expression (F_(1,20)_=33.86, p<0.0001, Figure 4F). We were unable to detect *SNAI2* transcript expression or vimentin protein expression in these cells, consistent with previous reports (63).

**Figure 4:**
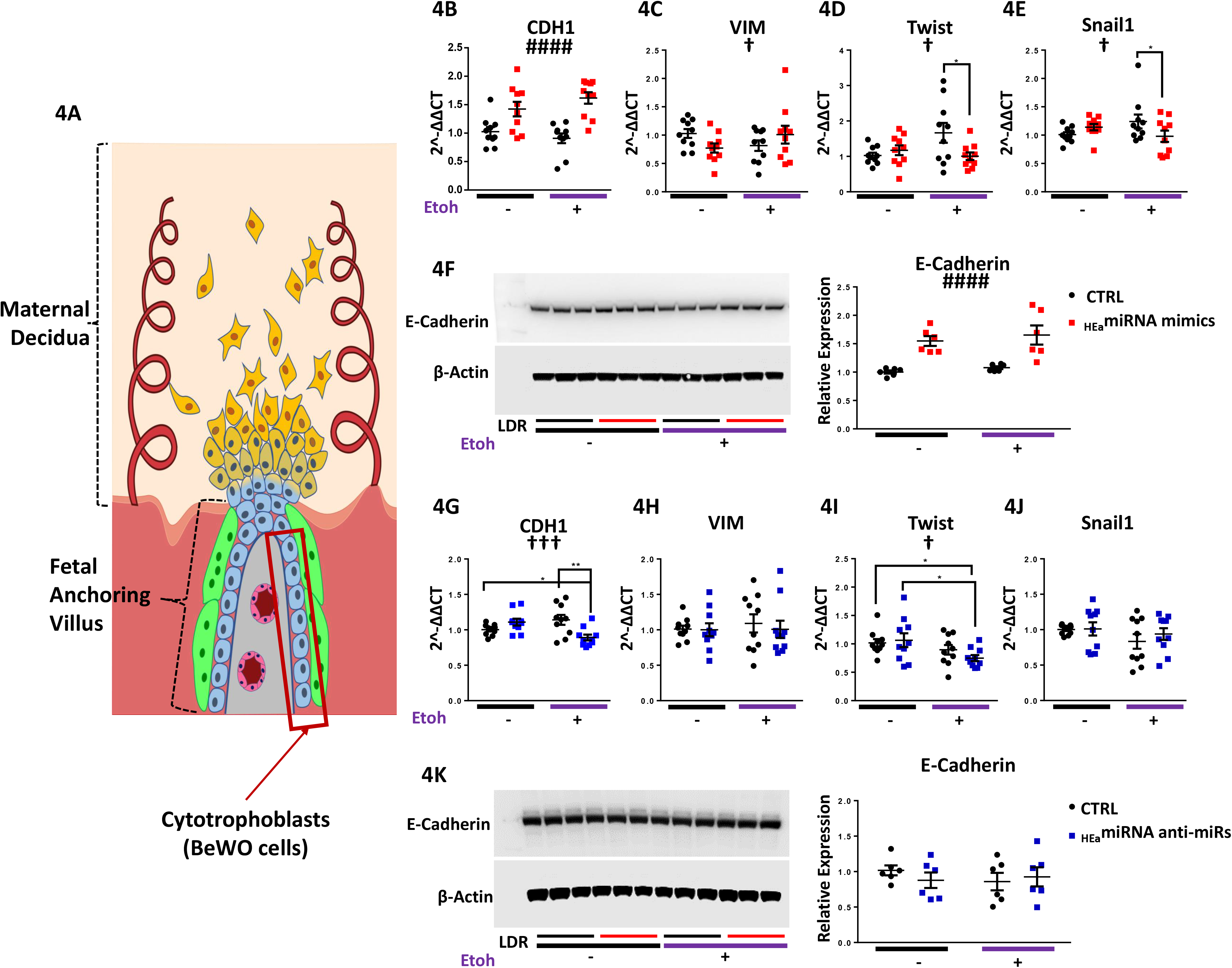
_HEa_miRNAs interfere with the EMT pathway in BeWO cytotrophoblasts. **A)** Diagram of a placental anchoring villous and maternal decidua with the boxed area denoting cytotrophoblasts. Expression of **B)** *CDH1*, **C)** *VIM*, **D)** *TWIST*, and **E)** *SNAI1* transcripts **F)** and densitometric quantification of E-Cadherin protein levels in BeWO cytotrophoblasts following _HEa_miRNAs or control miRNA overexpression with or without concomitant 320 mg/dL ethanol exposure. **G)** Expression of *CDH1*, **H)** *VIM*, **I)** *TWIST*, and **J)** *SNAI1* transcripts **K)** and densitometric quantification of E-Cadherin protein levels in BeWO cytotrophoblasts following _HEa_miRNAs or control hairpin inhibitor transfection with or without concomitant 320 mg/dL ethanol exposure. Results are expressed as the mean ± SEM, LDR=Molecular Weight Ladder, n=10 samples per group; ANOVA: significant main effect of _HEa_miRNA transfection [^####^p<0.0001], significant interaction effect (_HEa_miRNA by 320mg/dL ethanol, [^†^p<0.05, ^†††^p<0.001]). For post-hoc analysis *p<0.05, **p<0.01 by Tukey’s HSD.

We next sought to determine if more restricted subsets of _HEa_miRNAs could recapitulate the effects of _HEa_miRNAs collectively on EMT. Thus, we overexpressed hsa-miR-222-5p and hsa-miR-519a-3p, which are implicated in preeclampsia and fetal growth restriction, as well as hsa-miR-885-3p, hsa-miR-518f-3p, hsa-miR-204-5p, which are implicated in preeclampsia, fetal growth restriction, and spontaneous abortion or preterm labor (Supplementary Figure 6A). In contrast to the collective action for all _HEa_miRNAs, exposure to each of these pools resulted in significant decreases in CDH1 expression (F_(2,12)=_20.12, p=0.0001). The pool including hsa-miR-885-3p, hsa-miR-518f-3p, hsa-miR-204-5p also significantly increased Snai1 (F_(2,12)_=4.604, p=0.0328; Dunnett’s post-hoc p=0.0497, Supplementary Figure 6B-E). These data suggest that _HEa_miRNAs include sub-groups of miRNAs that have the potential to partly mitigate the effects of elevating the entire pool. However, the potential protective effects of these sub-groups are masked by the collective function of the entire group of _HEa_miRNAs.

Whereas transfection of _HEa_miRNA mimics increased *CDH1* expression, transfection of pooled antagomirs to _HEa_miRNAs, significantly reduced CDH1 expression, only in the context of 320 mg/dL ethanol co-exposure (_HEa_miRNA x 320mg/dL Etoh interaction, F_(1,36)_=13.51, p=0.0008; post-hoc Tukey’s HSD, p=0.005, Figure 4G). However, expression of *TWIST* was also decreased with ethanol co-exposure and there was no significant difference in E-Cadherin protein expression relative to the control (Figure 4H-K). Thus, our data suggest that, increasing _HEa_miRNA levels impairs EMT pathway members in cytotrophoblasts whereas inhibiting their action has a more restricted effect on EMT pathway members.

### _HEa_miRNAs impair EMT in a model of human extravillous trophoblasts

We next investigated the effect of _HEa_miRNAs on EMT in HTR-8/SVneo extravillous trophoblast-type cells (Figure 5A). Transfecting pooled _HEa_miRNA mimics into extravillous trophoblasts significantly decreased *VIM* expression (F_(1,36)_=28.43, p<0.0001). Expression of pro-mesenchymal transcription factors *SNAI2* was also reduced (F_(1,36)_= 64.88 respectively, p<0.0001). As with cytotrophoblasts, expression of SNAI1 and *TWIST* were reduced only with 320 mg/dL ethanol co-exposure (_HEa_miRNA x 320mg/dL Etoh interaction, F_(1,36)_=4.21 and 5.18, p=0.048 and 0.029 respectively; post-hoc Tukey’s HSD, p =0.027 and p<0.0001 respectively, Figures 5B-E). Consistent with our qPCR data, Vimentin protein expression was also significantly reduced (F_(1,20)_=9.535, p=0.006, Figure 5F). Interestingly, there was also a main effect of alcohol exposure on decreasing vimentin protein expression (F_(1,20)_=7.303, p=0.014). We were unable to detect expression of *CDH1* transcript, or its E-Cadherin protein product, in extravillous trophoblasts, consistent with previous reports (63).

**Figure 5:**
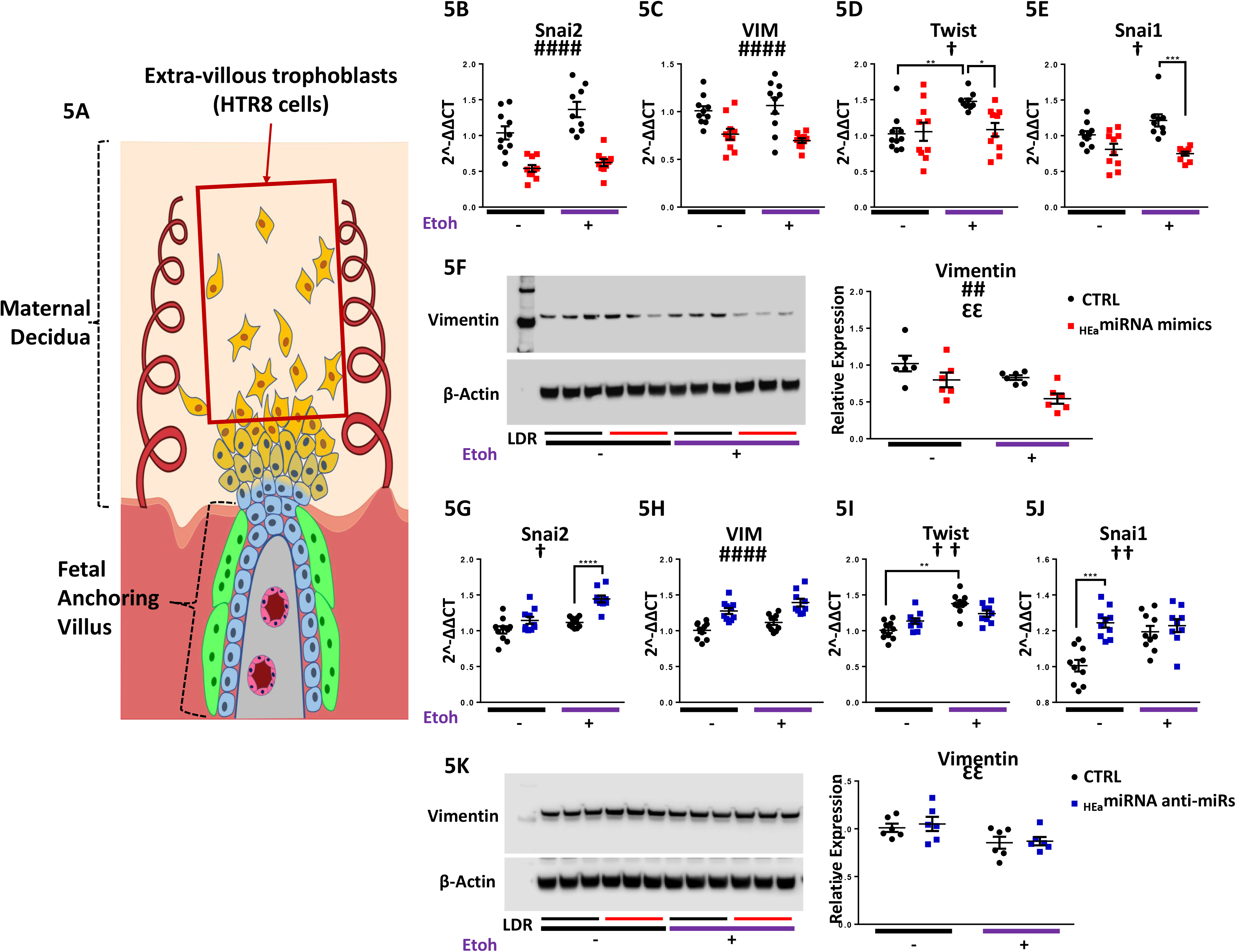
_HEa_mIRNAs Interfere with the IMT pathway In HTRB extravillous trophoblasts. **A)** Diagram of a placental anchoring villous and maternal decidua with the boxed area denoting extravillous trophoblasts. Expression of **B)** *SNAI2* **C)** *VIM* **D)** *TWIST* and **E)***SNAI1* transcripts **F)** as well as densitometric quantification of Vimentin protein levels in HTR8 extravillous trophoblasts following _HEa_rniRNAs or control miRNA overexpression with or with out concomitant 320 mg/dL ethanol exposure. Expression of **G)** *SNAI2* **H)** *VIM* **I)** *TWIST* and **J)** *SNAI1* transcripts **K)** as well as densitometric quantification of Vimentin protein levels in HTR8 extravillous trophoblasts following _HEa_miRNA or control hairpin inhibitor transfection with or without concomitant 320 mg/dL ethanol exposure. Results are expressed as the mean α SEM, LDR=Molecular WeiRht Ladder, n=10 samples per group; ANOVA: significant main effect of _HEa_miRNAtransfection [_##_p<0.01, _###_p<0.0001], significant main effect of 32Omg/dL ethanol exposure ⍰, significant interaction effect (_HEa_miRNA by 320mg/dLethanol, [^†^p<0.05, ^††^p<0.0l). For post hoc analysis *p0.05, **p<O.O1, ***p<0.001. and ***p<p<0. 0001 by Tukey’s HSD.

In contrast to _HEa_miRNA mimics, transfecting pooled antagomirs significantly increased *VIM* expression (F_(1,35)_=42.56, p<0.0001). Likewise, antagomir transfection increased expression of *SNAI2* in the context of 320mg/dL ethanol co-exposure and *SNAI1* under basal conditions (_HEa_miRNA x 320mg/dL Etoh interaction, F_(1,35)_=10.31 and 4.86, p=0.01 and p=0.034 respectively; post-hoc Tukey’s HSD, p<0.0001, Figures 5G-J). Despite our qPCR data, we did not observe significant differences in *VIM*entin protein expression between treatment groups (Figure 5K). Collectively, our data indicate that increased trophoblastic _HEa_miRNA levels favors an epithelial phenotype, whereas inhibiting their action promotes a mesenchymal phenotype.

### Antagomirs prevent _HEa_miRNAs’ inhibition of EMT

We next investigated if pretreating cytrophoblasts with pooled _HEa_miRNA antagomirs could prevent inhibition of the EMT pathway caused by transfecting _HEa_miRNA mimics. Pretreatment of cytotrophoblasts with _HEa_miRNA antagomirs prevented the elevation in *CDH1* caused by transfection with _HEa_miRNA mimics (post-hoc Tukey’s HSD, n=10 samples per group, p=0.004). Likewise, pre-transfection with _HEa_miRNA antagomirs also prevented _HEa_miRNA mimic induced reduction of *SNAI1* and *VIM* expression (post-hoc Tukey’s HSD, n=10 samples per group, p=0.007 and p<0.0001 respectively) (Figure 6A-D).

**Figure 6:**
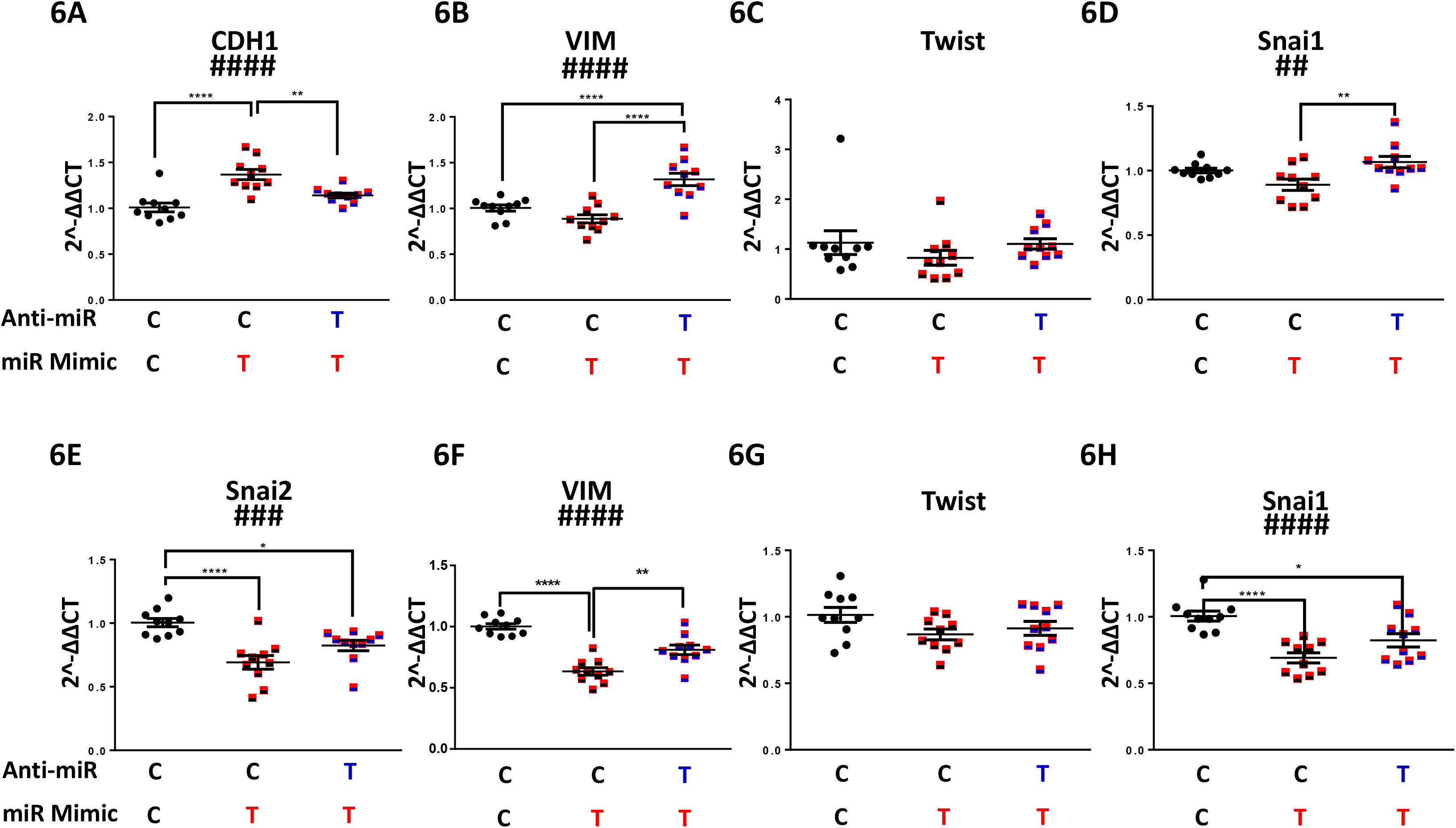
Antagomirs prevent _HEa_mIRNA induced impairment of EMT. Expression of **A)** *CDH1* **B)** *VIM* **C)** *TWIST* and **D)** *SNAI1* transcripts following control or _HEa_ miRNA hairpin inhibitor transfection followed by control or _HEa_ miRNA overexpression in BeWO cytotrophoblasts. Expression of **E)** *CDH1* **F)** *VIM* **G)** *TWIST* and **H)** *SNAI1* transcripts following control or _HEa_ miRNA antagomir transfection followed by control or _HEa_ miRNA overexpression in HTR8 extravillous trophoblasts. In subheadings: **C** denotes control miRNA mimic or hairpin whereas **T** denotes _HEa_ miRNA mimic or hairpin inhibitor. Results are expressed as expressed as the mean = SEM. n=10 samples per group; ANOVA: significant treatment effect [^##^ p<0.01, ^###^p<0.001. ^####^p<0.0001]. For post hoc analysis, *p<0.05, **p<0.01, ***p<0.001. **** p<0.0001 by Tukey’s HSD.

As with cytotrophoblasts, pre-transfection with _HEa_miRNA antagomirs prevented _HEa_miRNA mimic induced reduction of *VIM, SNAI1*, and *SNAI2* expression in extravillous trophoblasts (post-hoc Tukey’s HSD, n=10 samples per group, p<0.0001, Figure 6E-H). Thus, our data suggest that pretreating cells with _HEa_miRNA antagomirs prevents inhibition of EMT pathway members resulting from transfection with _HEa_miRNA mimics in cytotrophoblasts and extravillous trophoblasts.

### _HEa_miRNAs impair extravillous trophoblast invasion

Functionally, inhibition of the EMT pathway should reduce trophoblast invasiveness. Thus, we performed a transwell invasion assay using HTR8 extravillous trophoblasts transfected with _HEa_miRNA mimics and antagomirs. While ethanol exposure by itself did not impair trophoblast invasion (Supplementary Figure 7), there was a marginally significant interaction effect between ethanol exposure and _HEa_miRNA mimic transfection (F_(1,28)_=3.418, p=0.075). Thus, a planned comparison indicated that transfection with _HEa_miRNA mimics significantly reduced trophoblast invasion in the context of 320 mg/dL ethanol co-exposure, relative to the control mimics (t(14)=2.762, p=0.015), consistent with our data demonstrating _HEa_miRNAs interfere with the EMT pathway (Figure 7A). Contrastingly, transfecting _HEa_miRNA antagomirs increased invasion in the context of 320 mg/dL ethanol co-exposure, though this effect was only marginally significant (t(14)=1.805, p=0.093, Figure 7B).

**Figure 7:**
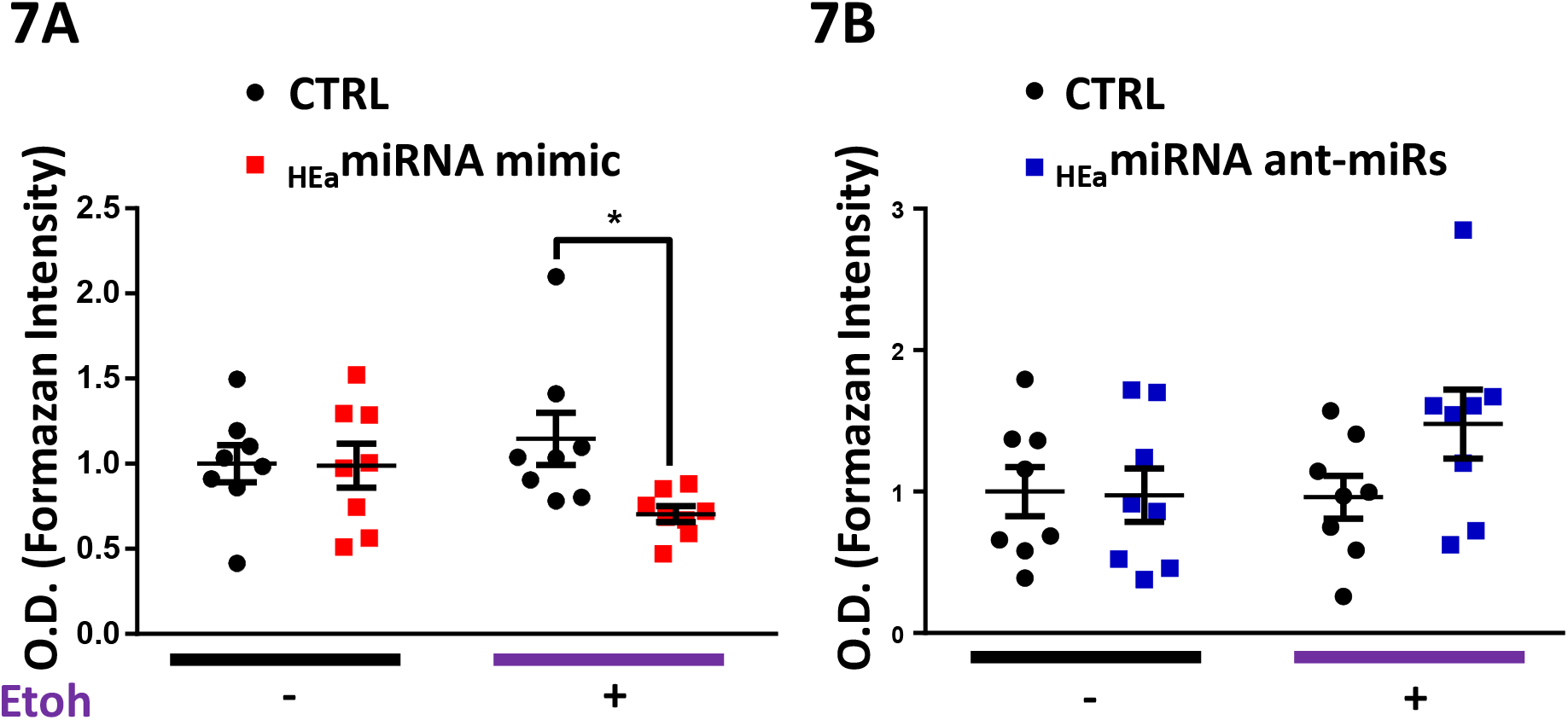
_HEa_ miRNA impair extravillous trophoblast invasion. Transwell invasion of HTR8 extravillous trophoblasts following transfection with **A) _HEa_** miRNA mimics or **B)** hairpin inhibitors with or without concomitant 320 mg/dL ethanol exposure. O.D. = optical density, results are expressed as expressed as the mean ± SEM; n=10 samples per group; *p<0.05 by Unpaired **T**-test

### _HEa_miRNAs retard trophoblast cell cycle progression

Given the proliferative nature of cytotrophoblasts, and the intimate relationship between EMT and cell cycle (64, 65), we assessed the effects of ethanol and _HEa_miRNAs on BeWO cytotrophoblast cell cycle. After pulse-labeling cells with the nucleic acid analog, EdU, for 1-hour, we found that individually transfecting 6 of the _HEa_miRNA mimics increased EdU incorporation (Unpaired t-test, p<0.05, FDR correction), suggesting an overall increased rate of DNA synthesis (Supplementary Figure 8A). Contrastingly, simultaneous transfection of _HEa_miRNAs significantly reduced EdU incorporation (F_(1,26)_=59.69, p<0.0001), mirroring the effects of increasing concentrations of ethanol (R^2^=0.304, p=0.012) (Supplementary Figure 8B and Figure 8A).

Consistent with the increased rates of DNA synthesis resulting from individual _HEa_miRNA mimic transfection, individual transfection of _HEa_miRNAs antagomirs generally reduced EdU incorporation, though only the antagomir to hsa-miR-760 did so significantly (t(110)=3.059, p=0.003, FDR correction) (Supplementary Figure 8A). Interestingly, simultaneous administration of antagomirs also reduced EdU incorporation, as observed with the pooled _HEa_miRNAs mimics (F_(1,26)_=34.83, p=0.0005, Figure 8B).

**Figure 8:**
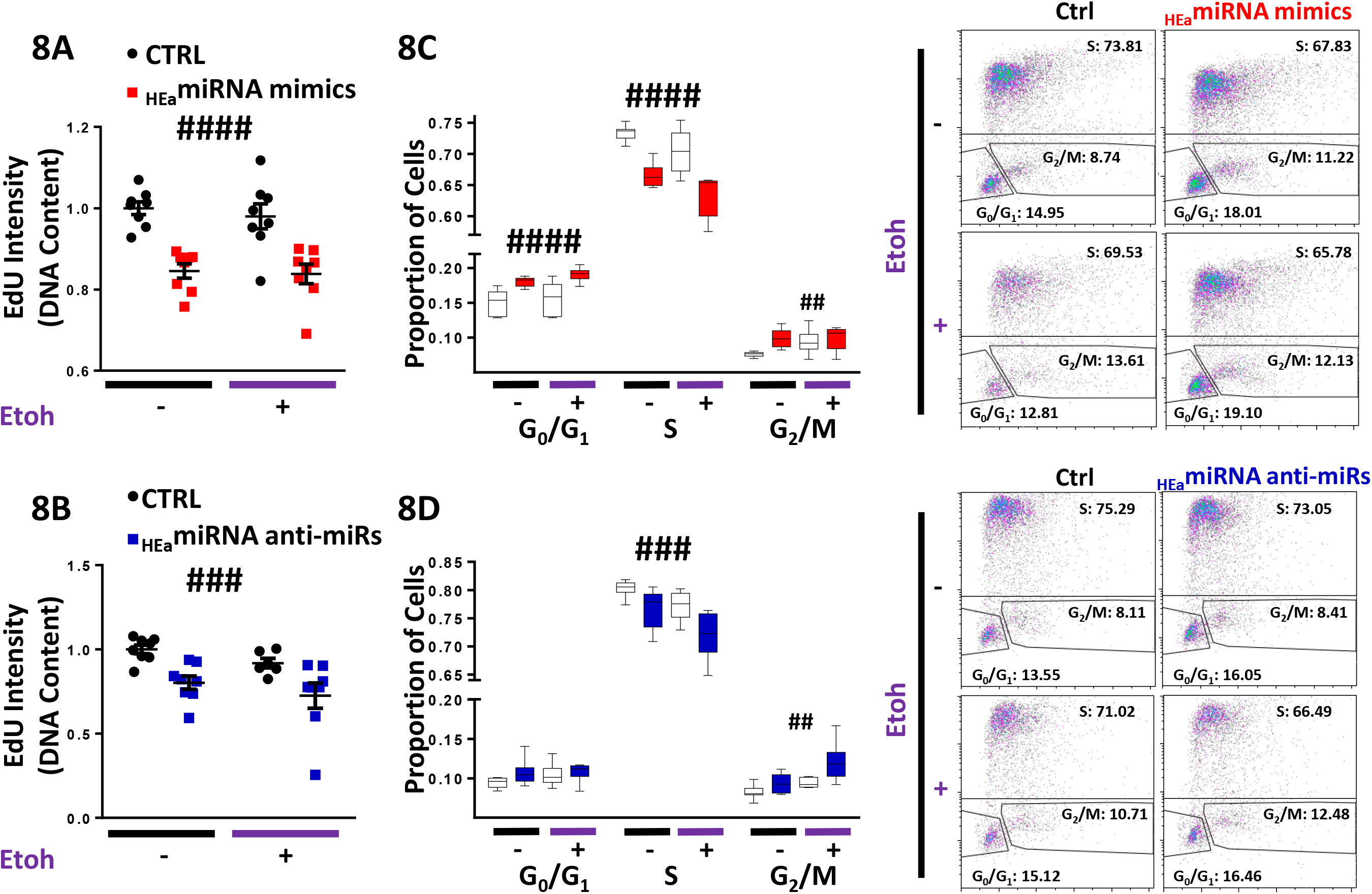
_HEa_ miRNA cause cell cycle retardation in trophoblasts. **A)** Degree of EdU incorporation following control and _Hea_ miRNA overexpression. **B)** Degree of EdU incorporation following control and _Hea_ miRNA hairpin inhibitor transfection. **C)** Box and whisker plot for the proportion of cells in G_0_ /G_1_, S, or G_2_ /M phase of the cell cycle following control and _HEa_ miRNA overexpression. **D)** Box and whisker plot for the proportion of cells in G_0_ /G_1_, S, or G_2_ /M phase of the cell cycle following control and _HEa_ miRNA hairpin inhibitor transfection with or without concomitant 320 mg/dL ethanol exposure. For box and whisker plots, bounds of box demarcate limits of 1st and 3rd quartile, line in middle is the median, and whiskers represent the range of data. Representative flow cytometry experiment images are shown on the right. n=10 samples per group; ANOVA: significant main effect of _HEa_miRNA transfection [^##^ p<0.01, ^###^p<0.001, and ^####^p<0.0001].

To further characterize the coordinated effect of _HEa_miRNAs on cytotrophoblast cell cycle, we pulse-labeled cells with EdU for 1-hour and, post-fixation, labelled them with 7AAD to segregate cells into three groups: G_0_/G_1_ (7AADlow, EDU-), S (EDU+), and G_2_/M (7AADhigh, EDU-). Both 120 mg/dL and 320 mg/dL ethanol exposures significantly decreased the proportion of cells in S-phase, while 320 mg/dL exposure increased the proportion of cells in G_2_/M-phase, consistent with the observed reduction in the rate of DNA synthesis (Supplementary Figure 8C). Similar, to the effects of ethanol exposure, pooled _HEa_miRNA mimic administration also significantly decreased the proportion of cells in S-phase (F_(1,28)_=52.78, p<0.0001) while increasing the proportion of cells the G_2_/M-phase (F_(1,28)_=8.395, p=0.007) and exacerbated alcohol’s effects on the cell cycle (Figure 8C). Interestingly, pooled _HEa_miRNA antagomir administration also reduced the proportion of cells in S-phase (F_(1,26)_=14.98, p=0.0007) and increased the proportion of those in G_2_/M-phase (F_(1,26)_=12.38, p=0.002) (Figure 8D).

As with our EMT gene expression data, pretreatment of cytotrophoblasts with antagomirs _HEa_miRNA prevented further reduction in the rate of DNA synthesis, or cell cycle retardation, that would result from transfection with pooled _HEa_miRNA mimics (Figures 9A and B).

**Figure 9:**
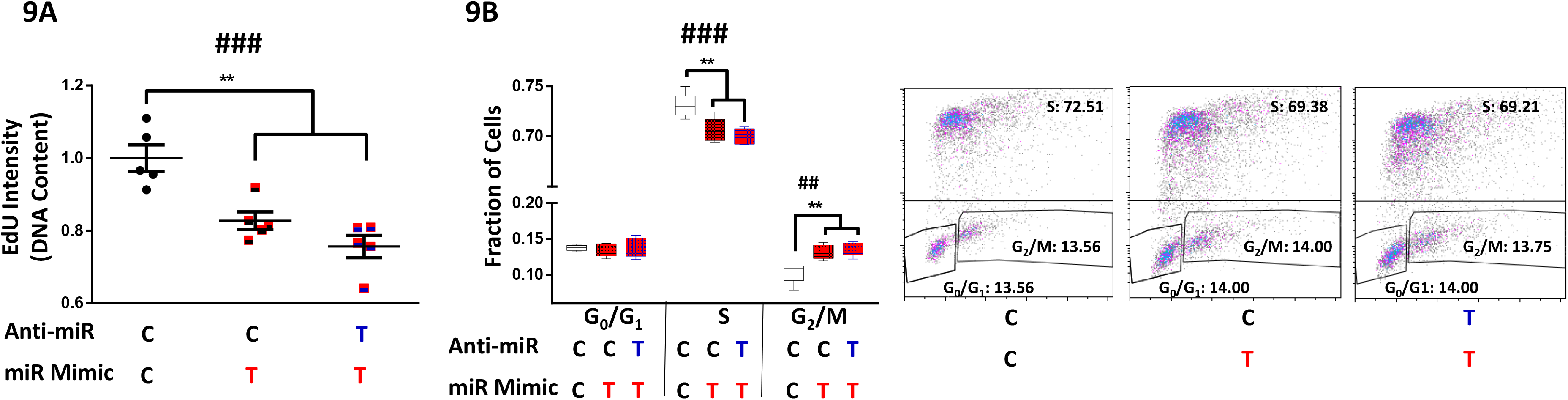
Antagomirs prevent _HEa_ miRNA induced cell cycle retardation. **A)** Degree of EdU incorporation following control or _HEa_ miRNA hairpin inhibitor transfection followed by control or _HEa_ miRNA overexpression in BeWO cytotrophoblasts. Results are expressed as expressed as the mean ± SEM. **B)** Box and whisker plot for the proportion of cells in G_0_ /G_1_, S, or G_2_ /M phase of the cell cycle following control or _HEa_ miRNA hairpin inhibitor transfection followed by control or _HEa_ miRNA overexpression in BeWO cytotrophoblasts. Bounds of box demarcate limits of 1st and 3rd quartile, line in middle is the median, and whiskers represent the range of data. Representative flow cytometry experiment images are shown on the right. In subheadings: **C** denotes control miRNA mimic or hairpin whereas T denotes miRNA mimic _Hea_ or hairpin inhibitor. n=5 samples per group; ANOVA: significant treatment effect [^###^ p<0.001]. For post-hoc analysis, **p<0.01 by Tukey’s HSD.

### _HEa_miRNAs have minimal effect on cell survival

We next investigated whether ethanol-and _HEa_miRNA-induced changes in cell cycle were related to an increase in cell death. Only the 320 mg/dL dose of ethanol exposure demonstrated a slight, but marginally significant effect, of increasing lytic cell death (t(18)=2.022, p=0.054), though there was no effect on apoptosis (Supplementary Figures 9A and B). However, the changes in cell cycle following transfection of individual or pooled _HEa_miRNA mimics were not mirrored by changes in lytic cell death. Nevertheless, two _HEa_miRNAs, hsa-mir-671-5p and hsa-mir-449a, did significantly increase apoptosis (Unpaired t-test, p<0.05, FDR correction) (Supplementary Figures 9C and D).

Contrastingly, transfection of 4 _HEa_miRNA antagomirs individually, significantly increased lytic cell death (Unpaired t-test, all p<0.05, FDR correction), with the antagomir to hsa-mir-491-3p also increasing apoptotic cell death (t(14)=3.383, p=0.004, FDR correction, Supplementary Figure 9C and D). Likewise, transfection of pooled _HEa_miRNA antagomirs increased lytic cell death (F_(1,36)_=11.40, p=0.002) but did not cause increased apoptosis (Supplementary Figure 9E-H). Taken together, our data suggest that while ethanol exposure may increase cytotrophoblast death, increased levels of _HEa_miRNAs have minimal effects on cell death, suggesting that their effect on cell cycle and the EMT pathway is independent of any effect on cell survival.

### _HEa_miRNAs modulate cytotrophoblast differentiation-associated Ca^2+^ dynamics

_HEa_miRNAs’ effects on EMT pathway member expression, coupled with cell cycle retardation, indicates that _HEa_miRNAs influence trophoblast maturation. To model _HEa_miRNAs’ effect on hormone-producing and calcium-transporting syncytiotrophoblasts (66), we used a well-established protocol of forskolin induced syncytialization of BeWO cytotrophoblasts (67, 68). As expected, forskolin treatment induced fusion/syncytialization of cytotrophoblasts resulting in a greater average cell size in the forskolin + _HEa_miRNA mimics group (F_(1,386)_=4.386, p=0.037). This suggests that the inhibition of EMT by these miRNAs may result in preferential syncytialization instead of differentiation to extravillous trophoblasts (Supplementary Figure 10A). Ethanol and forskolin treatment both increased baseline calcium levels, as indicated by the change in fluo-4 fluorescence (F_(1,426)_=5.593 and 3.665 respectively, p<0.0001, Figure 10A, Supplementary Figures 10B-D). The effect of ethanol on baseline calcium was abrogated by _HEa_miRNAs while _HEa_miRNAs + forskolin was not significantly different to forskolin alone, indicating that forskolin and _HEa_miRNAs may be affecting similar calcium pathways. The conversion of cytrophoblasts to syncytiotrophoblasts is accompanied by an increase in endoplasmic reticulum, which could increase calcium buffering capabilities in response to ethanol-stress on the cells, thus _HEa_miRNA-induced syncytialization pathways may be protective against ethanol stress.

**Figure 10:**
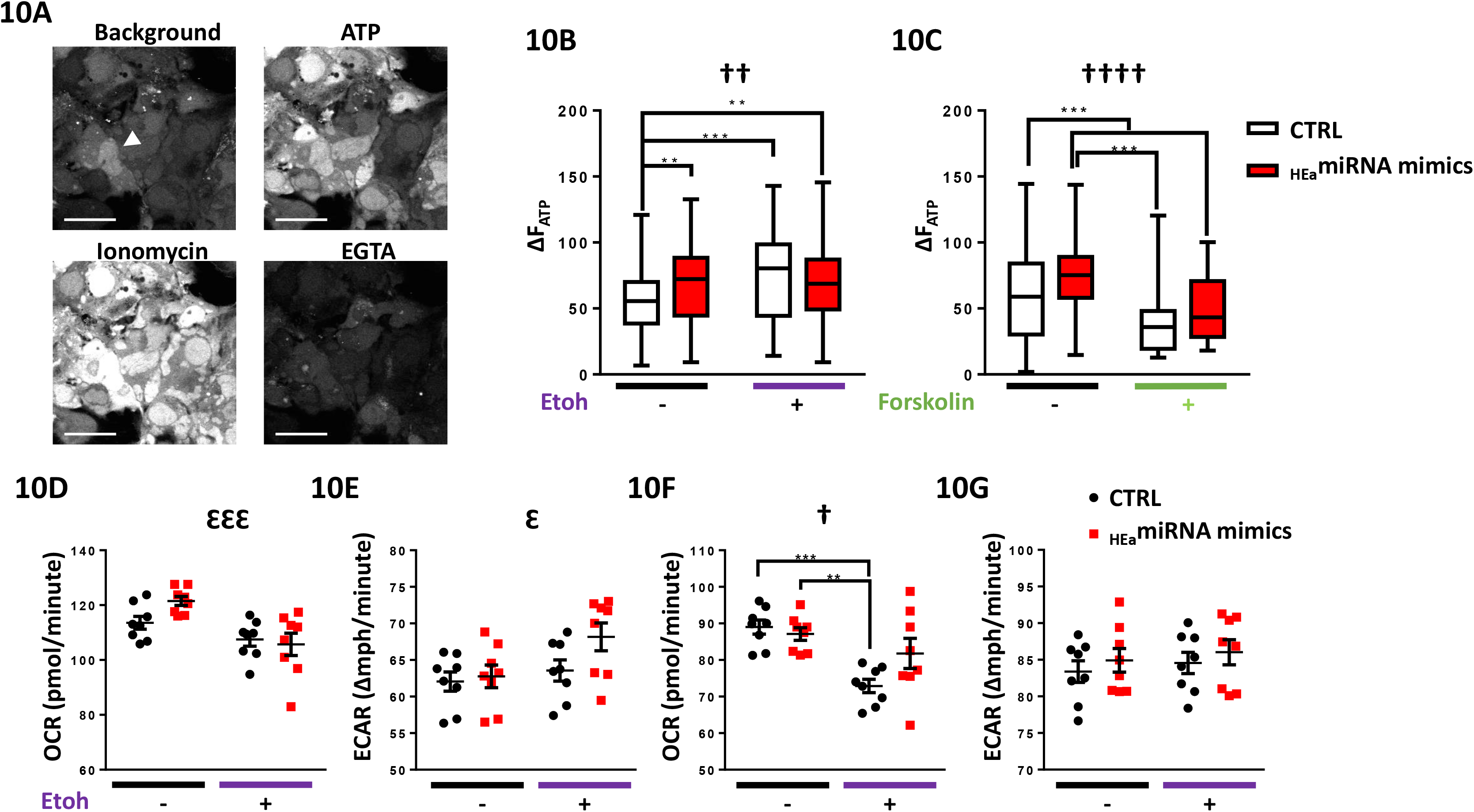
_Hea_ miRNAs modulate differentiation-associated Ca ^2+^ dynamics but have minimal effect on the cellular energetics profile. **A)** Time-lapse confocal images of BeWO cytotrophoblasts loaded with fluo-4 Ca **^2+^** indicator dye under indicated treatment conditions. Arrowhead indicates a fused, multinuclear cell, scale bar is 50µm. **B)** Box and whisker plot of intracellular calcium levels following acute ATP administration in BeWO cytotrophoblasts with control and _HEa_ miRNA overexpression with or without concomitant 320 mg/dL ethanol exposure. Bounds of box demarcate limits of 1^st^ and 3^rd^ quartile, line in middle is the median, and whiskers represent the range of data. **C)** Box and whisker plot of intracellular calcium levels following acute ATP administration in BeWO cytotrophoblasts with control and _HEa_miRNA overexpression with or without 20 μm forskolin treatment. **D)** Baseline oxygen consumption rate (OCR), **E)** baseline extracellular acidification rate (ECAR), **F)** stressed OCR, and 10G) stressed ECAR in BeWO cytotrophoblasts with control and _HEa_miRNA overexpression with or without concomitant 320mg/dL ethanol exposure. Metabolic stress was induced by treatment with 1μm Oligomycin and 0.125µM (FCCP). Results are expressed as expressed as the mean ± SEM. n=10 samples per group; ANOVA: significant main effect of 320mg/dL ethanol exposure 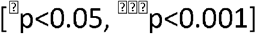, significant interaction effect (_HEa_miRNA by 320mg/dL ethanol, [^†^p<0.05, _††_p<0.01, and ^††††^p<0.0001]). For post-hoc analysis, *p<0.05, **p<0.01, ***p<0.001, and ***p<0.0001 by Tukey’s HSD.

Adaptations to cellular stress can also be seen in alterations to cellular energetics in response to ethanol, as ethanol-exposed BeWO cells showed decreased baseline and stressed oxygen consumption rates (OCR) (F_(1,28)_=15.55 and 16.91, p=0.0005 and 0.0003 respectively) and increased extracellular acidification rates (ECAR) (F_(1,28)_=4.868, p=0.036). However, _HEa_miRNAs had minimal effects on metabolic activity (Figures 10D-10G).

Extracellular ATP has been shown to inhibit trophoblast migration (69) and can directly stimulate increased intracellular calcium elevations through purinergic receptors ubiquitously present on trophoblasts (70). Both _HEa_miRNA and ethanol administration significantly increased intracellular calcium in response to acute ATP administration (F_(1,426)_=10.34 and F_(1,386)_=16.30, p=0.001 and p<0.0001 respectively) (Figure 10B). This may be indicative of a lack of downregulation of purinergic receptors required in trophoblast migration as part of the interrupted EMT pathway. Forskolin-induced maturation decreased calcium response to ATP (F_(1,386)_=50.72, p<0.0001) (Figure 10C) and prevented the _HEa_miRNA-induced increase in ATP response. These data agree with previous studies showing increased nuclear trafficking of ionotropic receptor P2X7 and more localized P2X4 expression over placental development, which may decrease the overall calcium influx in response to ATP (71).

### _HEa_miRNAs promotes syncytialization-dependent hormone production

Transfection of _HEa_miRNA mimics did not change *CGA, CGB*, or *IGF2* transcript expression relative to the control in non-syncytialized trophoblasts. However, following forskolin induced syncytialization of BeWO cytotrophoblasts (Figure 11A), _HEa_miRNA mimics significantly increased expression of *CGA* and *CGB* (post-hoc Tukey’s HSD, n=10 samples per group, p=0.001 and 0.005 respectively). Consistent with our previous results, _HEa_miRNA mimics also increased *CDH1* expression in both cytotrophoblasts and syncytiotrophoblasts (F_(1,20)_=5.286, p=0.032); there was also a main effect of syncytialization on *CDH1* expression, as has been previously reported (F_(1,36)_=3.391, p=0.034, Figures 11B-E). Likewise, _HEa_miRNAs increased E-cadherin protein expression (F_(1,20)_=5.286, p=0.032), whereas forskolin decreased it (F_(1,20)_=10.24, p=0.005) (Figure 11F). On the other hand, there was no effect of _HEa_miRNA antagomirs on *CGA* and *CGB* expression, although we did observe a decrease in *IGF2* transcript expression, following syncytialization, relative to controls (post-hoc Tukey’s HSD, n=10 samples per group, p=0.001) (Figure 11G-J).

**Figure 11:**
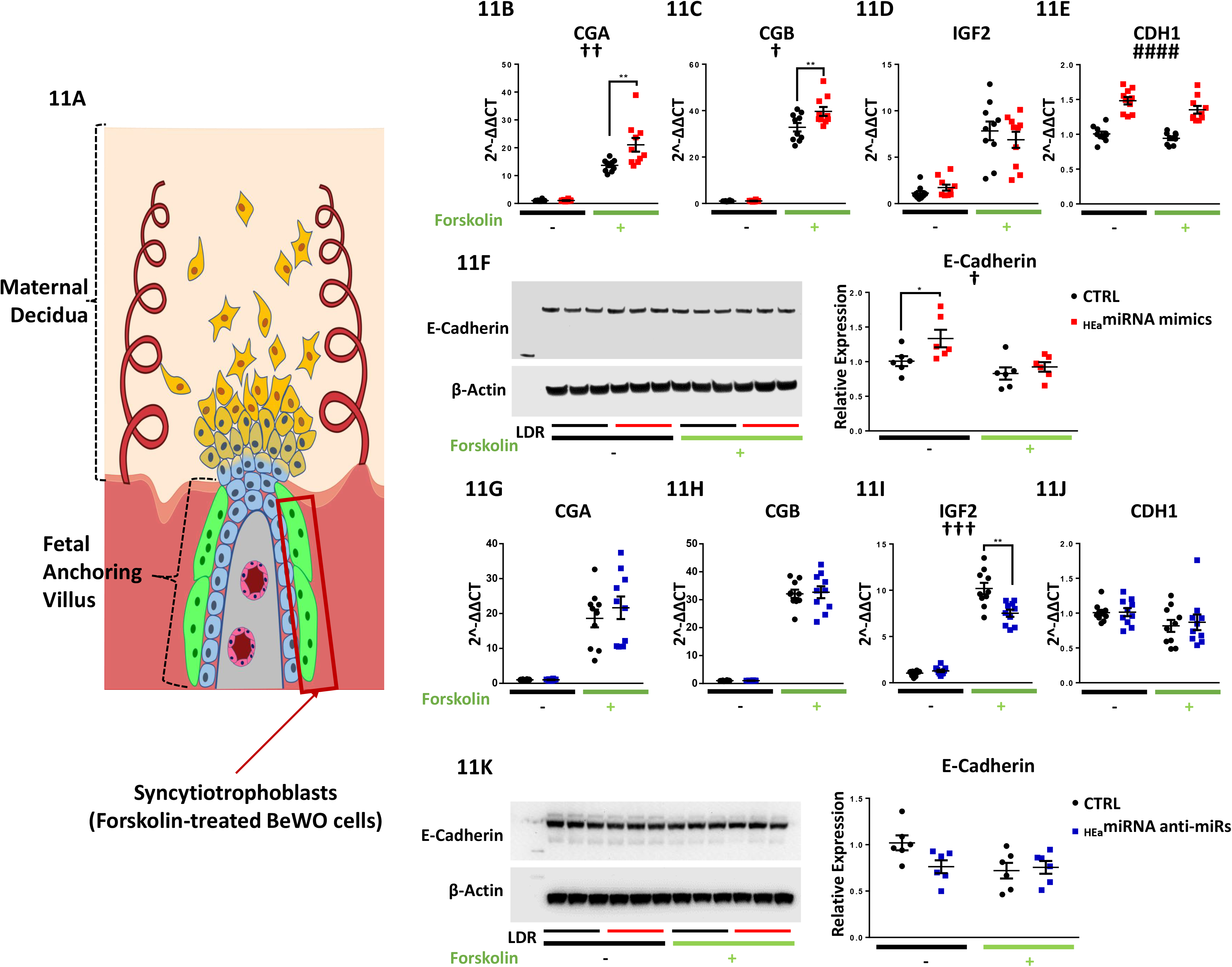
_HEa_miRNAs promote syncytialization dependent hCG production. **A)** Diagram of a placental anchoring villous and maternal decidua with the boxed area denoting syncytiotrophoblasts. Expression of **B)** *CGA*, **C)** *CGB*, **D)** *IGF2*, and **E)** *IGF2* transcripts **F)** and densitometric quantification of E-Cadherin protein levels in BeWO cytotrophoblasts following _HEa_miRNAs or control miRNA overexpression with or without 20 μm forskolin treatment. Expression of **G)** *CGA*, **H)** *CGB*, **I)** *IGF2*, and **J)** *IGF2* transcripts **K)** and densitometric quantification of E-Cadherin protein levels in BeWO cytotrophoblasts following _HEa_miRNAs or control hairpin inhibitor transfection with or without 20 μm forskolin treatment. Results are expressed as expressed as the mean ± SEM, LDR=Molecular Weight Ladder, n=10 samples per group; ANOVA: significant main effect of _HEa_miRNA transfection [_####_p<0.0001], significant interaction effect (_HEa_miRNA by 320mg/dL ethanol, [^†^p<0.05]). For post-hoc analysis, *p<0.05, **p<0.01 by Tukey’s HSD.

Given that _HEa_miRNAs promotes syncytialization-dependent hormone production, we next investigated maternal plasma levels of intact human chorionic gonadotropin (hCG) in our Ukraine birth cohort. Plasma hCG levels were non-significantly increased in the second trimester of HEa group mothers relative to their UE counterparts, consistent with previous studies (72). During the third trimester, however, hCG levels remained significantly elevated in HEa group mothers compared to the UE group (Median Test, n=23 samples in HEa group and n=22 for HEua and HEa groups, p=0.03) (Figure 12). Furthermore, there was no significant difference of gestational age at blood draw between the different groups indicating the increased level of hCG in the HEa group was not confounded by gestational age at which blood was sampled (Supplementary Figure 11) (73). Interestingly, both alcohol and hCG levels were negatively associated with gestational age at delivery (GAD), with a significant interaction between periconceptional alcohol exposure and hCG levels on GAD (Supplementary Table 3). Taken together, our data suggests _HEa_miRNAs may contribute to PAE-dependent increases in hCG levels during pregnancy.

**Figure 12:**
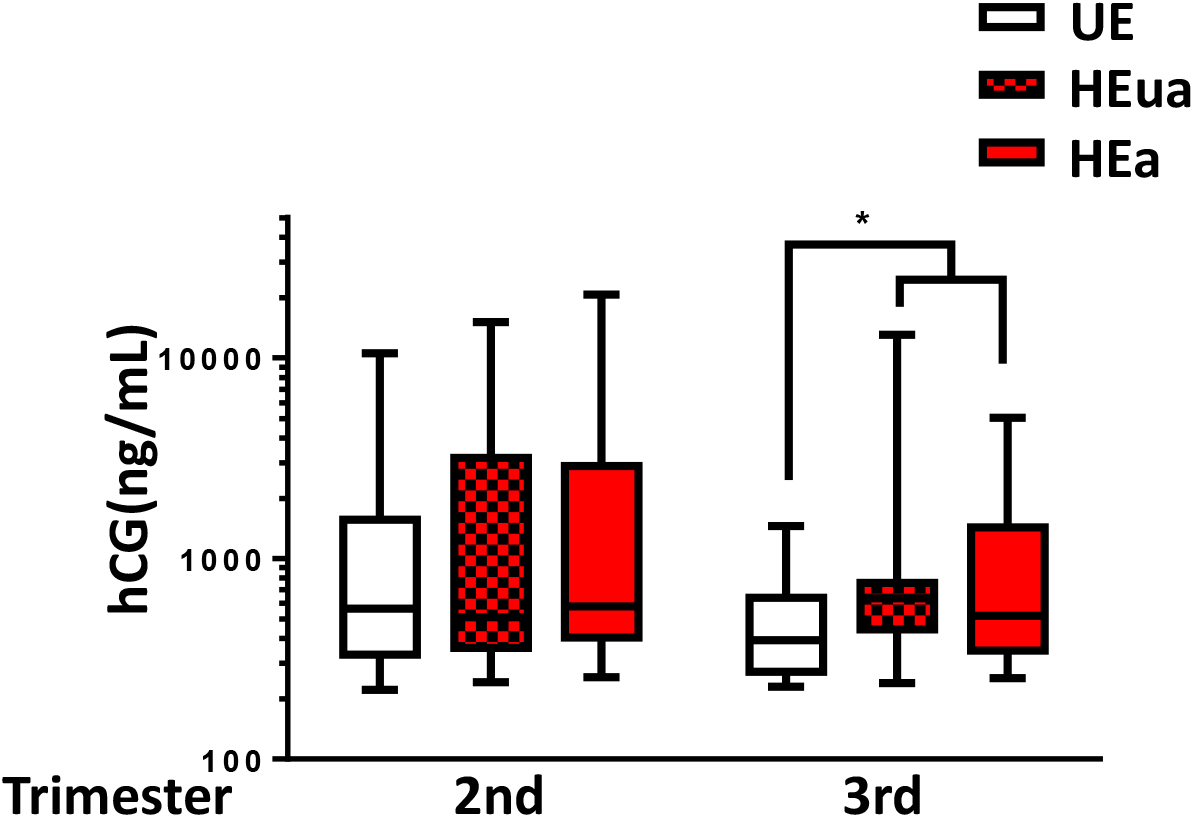
PAE elevates 3^rd^ trimester maternal hCG. Box and whisker plot of 2^nd^ and 3^rd^ trimester maternal hCG levels in UE, HEua, and HEa group mothers of our Ukrainian birth cohort. Bounds of box demarcate limits of 1st and 3rd quartile, line in middle is the median, and whiskers represent the range of data. Results are expressed as expressed as the mean ± SEM, n=22-23 samples per group; *p=0.03 (Mood’s Median Test, 𝒳_2_=7.043, df=2).

### _HEa_miRNAs reduce fetal growth

To investigate the functional consequences of elevated circulating _HEa_miRNA levels, we administered miRNA mimics for the 8-mouse homologue _HEa_miRNAs, or a negative control mimic, through tail-vein injection to pregnant mouse dams on GD10. On GD18, growth parameters of male and female fetuses were assessed separately, and data from all same-sex fetuses from a single pregnancy were averaged into one data point. Dams administered _HEa_miRNA mimics produced smaller fetuses than those administered control mimics, according to all collected measures of fetal size: fetal weight (F_(1,17)_=9.92, p=0.006), crown-rump length (F_(1,17)_=9.89, p=0.006), snout-occipital distance (F_(1,17)_=9.09, p=0.008), and biparietal diameter (F_(1,17)_=5.99, p=0.026) (Figure 13B-E). Interestingly, placental weights were also significantly reduced in mice treated with _HEa_miRNA mimics (F_(1,17)_=6.92, p=0.018) (Figure 13F).

**Figure 13:**
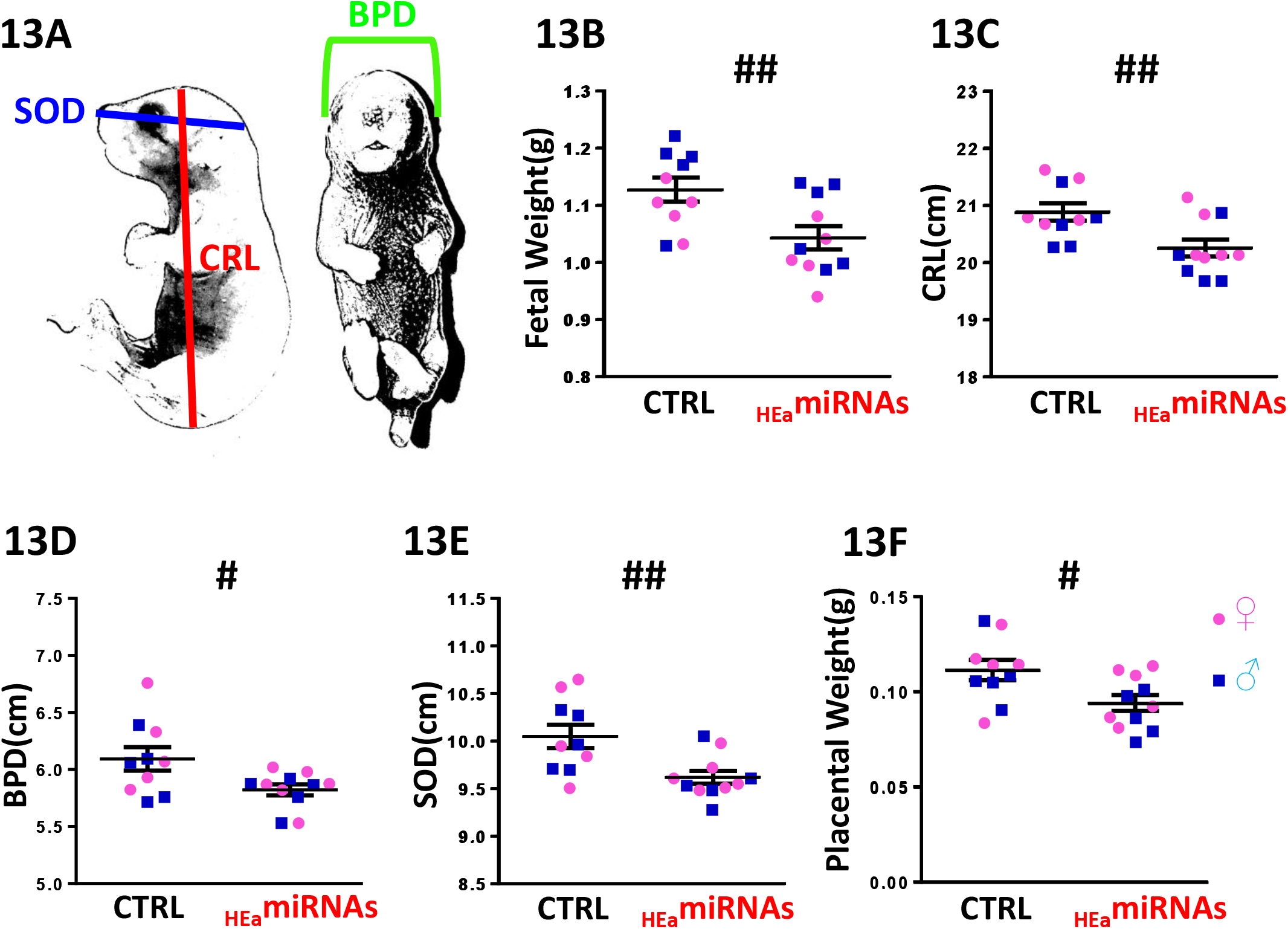
_HEa_miRNAs restrict fetal growth. **A)** Schematic for measures of crown rump length (CRL), biparietal diameter (BPD), and snout-occipital distance (SOD). **B)** Fetal weight, **C**) crown-rump length, **D)** biparietal diameter, **E)** snout-occipital distance, **F)** and placental weight at GD18 following administration of control (Ctrl) and _HEa_miRNA mimics to pregnant C57/Bl6 dams on GD10. Dots represent median measures of fetal size and placental weights from male and female offspring in independent litters. There were no significant differences in litter sizes [Ctrl: 8.2 and _HEa_miRNAs: 8.5] or sex ratios [Ctrl: 0.86 and _HEa_miRNAs: 1.21] between treatment conditions (p>0.5 for all measures). Results are expressed as expressed as the mean ± SEM, n=5-6 separate litters per treatment condition; ANOVA: significant main effect of _HEa_miRNA administration [_#_p<0.05 and _##_p<0.01].

Following tail-vein administration of two human-specific sentinel miRNAs, miR-518f-3p and miR-519a-3p, we found a high biodistribution of both miRNAs in the placenta, comparable to levels seen in the liver and spleen (Supplementary Figure 12A and 12B). Thus, to determine whether _HEa_miRNA’s effects on fetal growth could result from their actions on the placenta, we quantified the placental expression of core EMT members in the GD18 placentas of control and _HEa_miRNA fetuses. _HEa_miRNA administration significantly reduced expression of mesenchymal-associated transcript *VIM* (F_(1,14)_=14.23, p=0.002) and *SNAI2* (F_(1,14)_=5.99, p=0.028) with a significant sex by _HEa_miRNA interaction effect on *SNAI1* (F_(1,66)_=5.55, p=0.034) and *CDH1* (F_(1,14)_=6.01, p=0.028) (Figures 14A-E). Interestingly, and in line with our *in vitro* findings whereby _HEa_miRNAs promoted syncytialization dependent cell fusion and hCG production, _HEa_miRNA administration significantly increased expression of the mRNA transcript for *SynB*, a gene that is important for syncytiotrophoblast maturation (F_(1,66)_=4.11, p=0.047) (Figure 14F).

**Figure 14:**
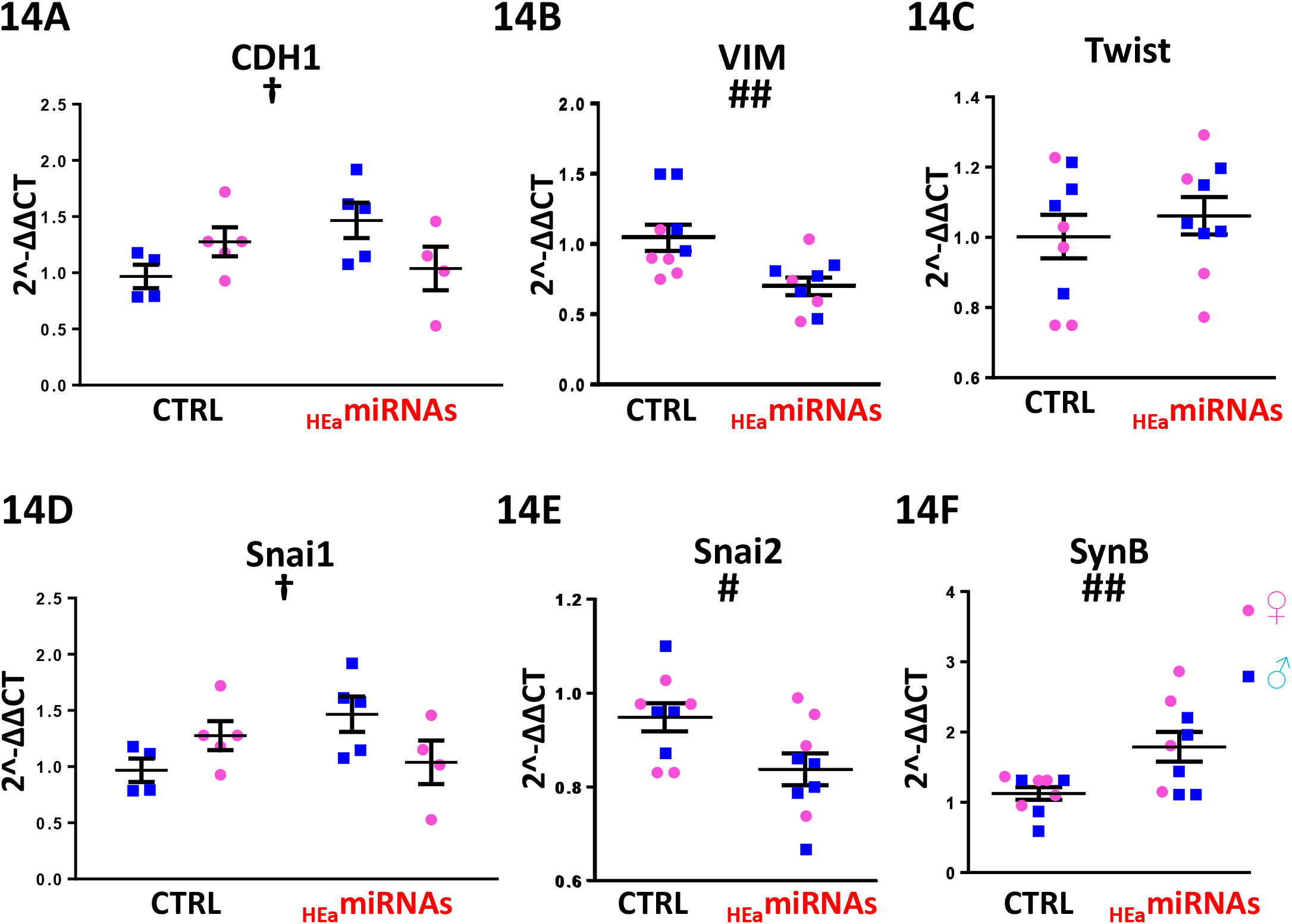
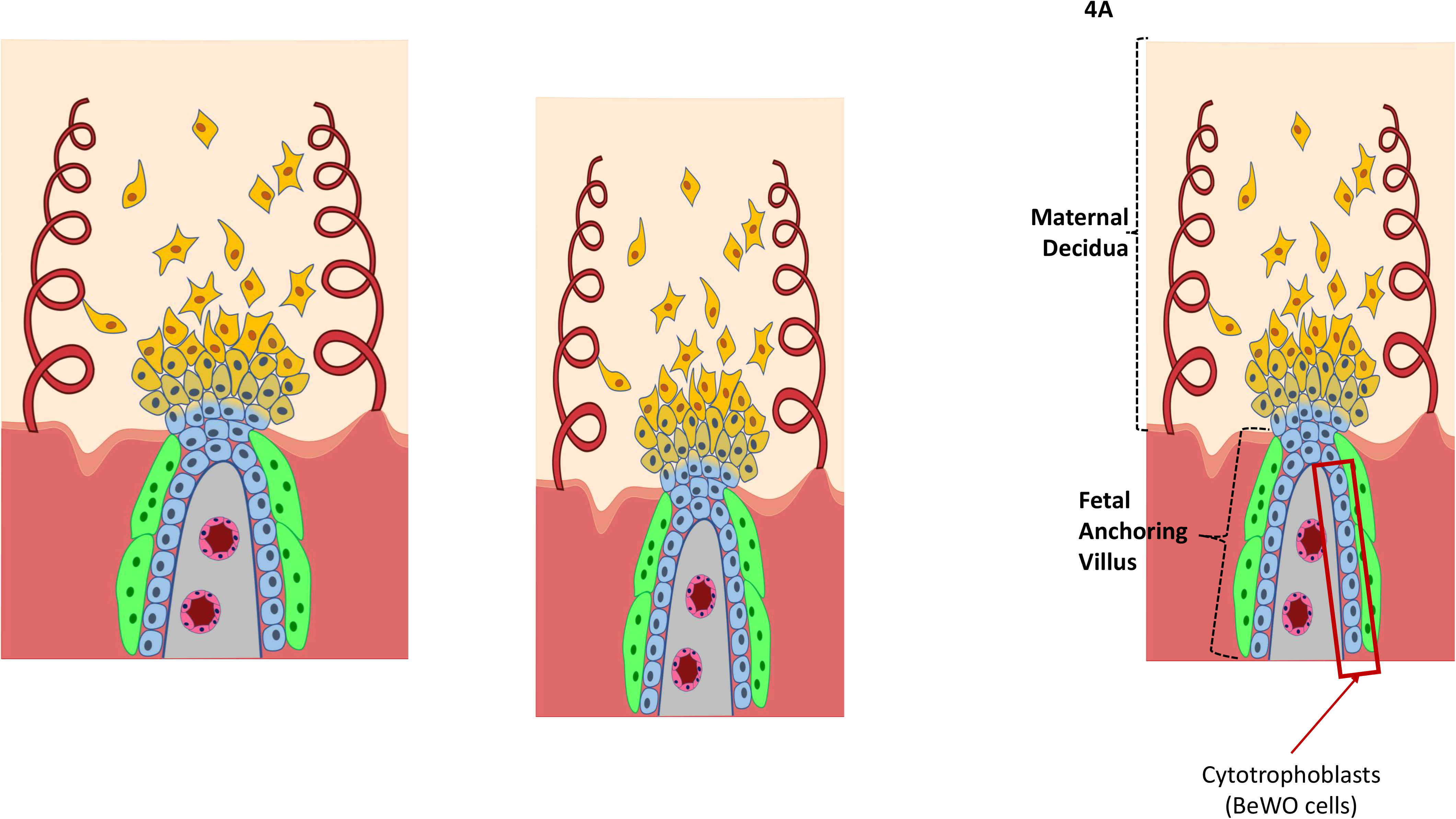
_HEa_miRNAs interfere with EMT in the placenta. Expression of **A)** *CDH1* **B)** *VIM* **C)** *TWIST* **D)** *SNAI1* and **E)** Snai2 and **F)** *SynB* transcripts in GD18 placenta following administration of control (Ctrl) and _HEa_miRNA mimics to pregnant C57/Bl6 dams on GD10. Dots represent median expression values of male and female offspring in independent litters. Results are expressed as expressed as the mean ± SEM, n=5-6 separate litters per treatment condition, ANOVA: significant main effect of _HEa_miRNA administration [_#_p<0.05, _###_p<0.001], significant interaction effect (fetal sex by _HEa_miRNA administration, [^†^p<0.05]). For post-hoc analysis, *p<0.05 by Tukey’s HSD.

## Discussion

We previously reported that gestational elevation of 11 maternal plasma miRNAs predicted which PAE infants would exhibit adverse outcomes at birth (8). These _HEa_miRNAs were elevated throughout mid and late-pregnancy, encompassing critical periods for fetal development, and were predicted to target the EMT pathway (8). In this study, we tested this prediction by adopting rodent and macaque gestational moderate alcohol self-administration paradigms. Despite differences in their placental anatomy (74–77), we are the first to report that PAE impairs placental EMT across species, indicating a conserved effect of PAE on placental development. Additionally, we found that _HEa_miRNAs collectively, but not individually, mediated the effects of PAE on core EMT pathway members and that, together, they inhibited EMT in human trophoblast culture models. While we assessed the effects of _HEa_miRNAs on core EMT components (10,14,15, 59–62), analysis of their 3’UTRs indicates that these are unlikely to be the direct targets of _HEa_miRNA action. Additional studies will be needed to dissect out the signaling networks that connect _HEa_miRNAs to the assessed EMT components.

Interestingly, _HEa_miRNAs also promoted syncytialization (forskolin)-dependent hCG expression, mirroring the elevation of third trimester maternal hCG levels in the PAE group within our clinical cohort. This late-gestation elevation of hCG levels may serve as a compensatory mechanism to prevent the preterm birth associated with PAE, as hCG during late gestation is hypothesized to promote uterine myometrial quiescence (78, 79). In support of this hypothesis, we found significant negative associations between both hCG levels and alcohol consumption with gestational age at delivery. Furthermore, there was a significant interaction between periconceptional alcohol exposure and hCG levels, with higher hCG levels corresponding to a smaller effect of alcohol exposure at conception on gestational age at delivery, indicating that hCG moderates the effect of alcohol on age at delivery (Supplementary Table 3).

Since _HEa_miRNAs collectively prevented trophoblast EMT, we hypothesized that, as a functional consequence, these maternal miRNAs would also inhibit fetal growth. When we delivered 8 out of the 11 _HEa_miRNAs known to be present in mouse, to pregnant dams during the period of placental branching morphogenesis and endometrial invasion, when EMT is particularly active, we found that _HEa_miRNAs reduced fetal growth. Importantly, ethanol exposure during this period has also been shown to result in fetal growth deficits and dysmorphia in rodent PAE models (80, 81) suggesting that maternal miRNA-mediated deficits in trophoblast invasion may mediate some of the effects of PAE on fetal growth. In support of this, we found placentas from the _HEa_miRNA treated group had impaired expression of core EMT pathway members. This disruption of placental EMT may also have implications for placental vascular dynamics, as we have also previously observed in mouse models (82). The non-human primate tissue analyzed here was also derived from animals that were characterized *in vivo* using MRI and ultrasound imaging, which demonstrated that maternal blood supply to the placenta was lower in ethanol-exposed animals compared to controls, and that oxygen availability to the fetal vasculature was reduced (83).

_HEa_miRNAs may mediate other pregnancy associated pathologies, aside from PAE. We identified numerous studies that reported increased circulating and placental levels of at least 8 out of 11 _HEa_miRNAs in gestational pathologies arising from placental dysfunction. For example, elevated levels of one _HEa_miRNA, miR-519a-3p, a member of the placentally-expressed C19MC family cluster, was reported in placentae of patients with pre-eclampsia, recurrent spontaneous abortion, and intrauterine growth restriction (29, 30, 45,46). Interestingly, collective overexpression of the 59 C19MC miRNAs inhibits trophoblast migration, explaining their enrichment in the non-migratory villous trophoblasts and suggests their downregulation is necessary for maturation into invasive extravillous trophoblasts (84). Thus, a greater understanding of the placental roles of _HEa_miRNAs may also help disentangle the etiology of other pregnancy complications. We also observed that overexpression of more restricted subsets of _HEa_miRNAs associated with preeclampsia, fetal growth restriction, and spontaneous abortion or preterm labor also partly promoted EMT transcript signatures, contrasting with the collective inhibitory action of _HEa_miRNAs as a whole. Thus, elevation of some subsets of _HEa_miRNAs may constitute a compensatory mechanism aimed at minimizing placental pathologies, though their potential protective effects are masked by the collective elevation of _HEa_miRNAs.

While we did not investigate the effects of PAE on EMT in non-placental organs, it is likely that PAE broadly disrupts EMT in multiple fetal compartments. Developmental ethanol exposure has been shown inhibit the EMT-dependent migration of neural crest progenitors involved in craniofacial development, explaining the facial dysmorphology seen in FAS and FASDs (85, 86). Outside of its effects on the neural crest, PAE is significantly associated with various congenital heart defects, including both septal defects and valvular malformations (87–90). Given that development of heart depends on EMT within the endocardial cushions (91, 92), disruption of endocardial EMT could explain both the valvular and septal malformation associated with PAE.

Collectively, our data on _HEa_miRNAs suggest miRNA-based interventions could minimize or reverse developmental effects of PAE and other placental-related pathologies. miRNA-based therapeutic approaches have been advanced for other disease conditions(93)(94). However, our data also suggests the effects of combinations of miRNAs are not a sum of their individual effects. Functional synergy between clusters of co-regulated miRNAs may be a common feature in development and disease. For instance, in 2007, we presented early evidence that ethanol exposure reduced miR-335, −21, and -153 in neural progenitors and that coordinate reduction in these miRNAs yielded net resistance to apoptosis following ethanol exposure (95). In that study, we also showed that coordinate knockdown of these three miRNAs was required to induce mRNA for Jagged-1, a ligand for the Notch cell signaling pathway, an outcome that was not recapitulated by knocking down each miRNA individually (95). More recently, combined administration of miR-21 and miR-146a has been shown to be more effective in preserving cardiac function following myocardial infarction than administration of either of these miRNAs alone (96). While miRNA synergy has not been explored in detail, these data show that new biology may emerge with admixtures of miRNAs, and that therapeutic interventions may require the use of such miRNA admixtures rather than single miRNA molecules, as have been used in clinical studies to date.

In conclusion, we have observed that a set of 11 miRNAs, predictive of adverse infant outcomes following PAE, collectively mediate the effects of alcohol on the placenta. Specifically, elevated levels of these miRNAs together, but not individually, promote an aberrant maturational phenotype in trophoblasts by inhibiting core members of the EMT pathway and promoting cell stress and syncytialization-dependent hormone production. While extensive research has established circulating miRNAs as biomarkers of disease, our study is one of the first to show how these miRNAs explain and control the disease process themselves. Functionally, we find that these miRNAs are clinically correlated with measures of fetal development and directly cause intrauterine growth restriction when administered *in vivo.* Our work suggests that a greater understanding for the role of _HEa_miRNAs during development, and their role in coordinating the EMT pathway in the placenta and other developing tissues, will benefit the understanding of FASDs and other gestational pathologies and potentially lead to effective avenues for intervention.

## Methods

### Mouse model of PAE

C57/BL6J mice (Jackson Laboratory, Bar Habor, ME) were housed under reverse 12-hour dark / 12-hour light cycle (lights off at 08:00 hours). PAE was performed using a previously described limited access paradigm of maternal drinking (97, 98). Briefly, 60-day old female mice were subjected to a ramp-up period with 0.066% saccharin containing 0% ethanol (2 days), 5% ethanol (2 days), and finally 10% ethanol for 4-hours daily from 10:00–14:00 beginning 2 weeks prior to pregnancy, continuing through gestation (Supplementary Figure 2A). Female mice offered 0.066% saccharin without ethanol during the same time-period throughout pregnancy served as controls. Tissue from the labyrinth, junctional, and decidual zone of male and female gestational day 14 (GD14) placentae were microdissected, snap-frozen in liquid nitrogen, and stored at −80⍰C preceding RNA and protein isolation.

### Mouse model for _HEa_miRNA overexpression

For systemic administration of miRNAs, previously nulliparous C57/BL6NHsd dams (Envigo, Houston, TX) were tail-vein-injected on GD10 with either 50 μg of miRNA miRVana™ mimic negative control (Thermo Fisher, Waltham, MA, Cat No. 4464061) or pooled _HEa_miRNA miRVana™ mimics in In-vivo RNA-LANCEr II (Bioo Scientific, Austin, TX, 3410-01), according to manufacturer instructions. The 50 μg of pooled _HEa_miRNA mimics consisted of equimolar quantities of mmu-miR-222-5p, mmu-miR-187-5p, mmu-mir-299a, mmu-miR-491-3p, miR-760-3p, mmu-miR-671-3p, mmu-miR-449a-5p, and mmu-miR-204-5p mimics. For bio-distribution studies, 50 μg of pooled equimolar quantities of hsa-miR-519a-3p and hsa-miR-518f-3p mimics were injected via tail vein. These human miRNAs were selected because no mouse homologs are known to exist and consequently, estimates for organ distribution of exogenous miRNAs in the mouse are unlikely to be contaminated by the expression of endogenous murine miRNAs. GD10 is a time point near the beginning of the developmental period of branching morphogenesis, immediately following chorioallantoic attachment, during which the placenta invades the maternal endometrium (99). At GD18, pregnancies were terminated with subsequent quantification of fetal weight, crown-rump length, snout-occipital distance, biparietal diameter, and placental weight (Figure 13A). Subsequently, tissue was snap-frozen in liquid nitrogen, and stored at ×80⍰C preceding RNA isolation.

### Rat model of PAE

Outbred nulliparous Sprague-Dawley rats were housed under a 12-hour light/12-hourdark cycle. PAE in Sprague-Dawley was conducted according to our previously published exposure paradigm (20, 100). Briefly, dams were given a liquid diet containing either 0% or 12.5% ethanol (vol/vol) from 4 days prior to mating until GD4 (Supplementary Figure 2B). Dams had ad libitum access to the liquid diet 21-hours daily and consumed equivalent calories. Water offered during the remaining 3-hours of the day. On GD5, liquid diets were removed and replaced with standard laboratory chow. On GD20, placentas were immediately separated into the labyrinth and junctional zone, snap frozen in liquid nitrogen and stored at −80 °C preceding RNA isolation.

### Non-human primate model of PAE

As previously described in detail (83), adult female rhesus macaques were trained to orally self-administer either 1.5 g/kg/d of 4% ethanol solution (equivalent to 6 drinks/day), or an isocaloric control fluid prior to time-mated breeding. Each pregnant animal continued ethanol exposure until gestational day 60 (GD60, term gestation is 168 days in the rhesus macaque) (101). Pregnancies were terminated by cesarean section delivery at three different time points; GD85, GD110, or GD135 (Supplementary Figure 2C). The macaque placenta is typically bi-lobed with the umbilical cord insertion in the primary lobe and bridging vessels supplying the fetal side vasculature to the secondary lobe (Figure 2D showing gross placenta anatomy) (102). Full thickness tissue biopsies (maternal decidua to fetal membranes) were taken from both the primary and secondary lobes of the placenta (Figure 2E showing H&E section of placenta). Samples were immediately snap-frozen in liquid nitrogen and stored at ×80°C preceding RNA isolation.

### Cell culture trophoblast models

BeWO human cytotrophoblastic choriocarcinoma cells and HTR-8/SVneo extravillous cells were sourced from ATCC (Manassas, VA, Cat No. CCL-98 and CRL-3271 respectively). BeWO cells were maintained in HAM’s F12 media containing penicillin (100 U/ml), streptomycin (100 μg/ml), and 10% vol/vol fetal calf serum (FCS) at 37°C and 5% CO_2_. HTR8 cells were maintained in RPMI-1640 media with 5% vol/vol FCS, under otherwise identical conditions. Culture medium was replenished every 2 days and cells sub-cultured every 4-5 days.

BeWO cells were treated with 20 µm forskolin to induce syncytialization, as previously described (103, 104). BeWO and HTR8 cells were also subjected to four separate ethanol treatment conditions: 0 mg/dL, 60 mg/dl (13 mM),120 mg/dl (26 mM) or 320 mg/dl (70 mM). To achieve _HEa_miRNA overexpression and inhibition, Dharmacon miRIDIAN™ miRNA mimics and hairpin inhibitors [25 nM], or control mimic (Dharmacon, Lafeyette CO, Cat No. CN-001000-01-05) and hairpin inhibitor (Dharmacon, Cat No. CN-001000-01-05) [25nm], were transfected into subconfluent BeWO and HTR8 cells using RNAIMAX lipofection reagent (Thermo Fisher, Cat No. 13778).

### Cell cycle analysis

At 48-hours-post transfection, BeWO cells were pulsed with 10 μM EdU for 1-hour. Cells were immediately harvested, and cell cycle analysis was performed with the Click-iT® EdU Alexa Fluor® 488 Flow Cytometry Assay Kit (Thermo Fisher, Cat No. C10420), in conjunction with 7-Amino-Actinomycin D (Thermo Fisher, Cat No. 00-6993-50), according to manufacturer instructions, using the Beckman Coulter® Gallios 2/5/3 Flow Cytometer. Data was analyzed using Kaluza software (Beckman Coulter, Brea, CA).

### Cell death analysis

BeWO cell culture was harvested 48-hours post transfection media was subjected to lactate dehydrogenase (LDH) detection using the Pierce™ LDH Cytotoxicity Assay Kit (Thermo Fisher, Cat No. 88953), according to manufacturer instructions, for lytic cell death quantification. The Promega Caspase-Glo® 3/7 Assay Systems (Promega, Madison, WI, Cat No. G8091) was used to quantify apoptotic cell death

### Invasion assay

At 24-hours post-transfection and/or ethanol exposure, HTR8 cells were serum starved for an additional 18-hours. Subsequently, HTR8 cells were seeded onto trans-well permeable supports precoated with 300 µg/mL Matrigel (Corning, Corning, NY, Cat No. 354248). After 24-hours, cells remaining in the apical chamber were removed with a cotton swab. Cells that invaded into the basal chamber were incubated with 1.2 mM 3-(4,5-dimethylthiazol-2-yl)-2,5-diphenyltetrazolium bromide (MTT) for 3-hours, and the precipitate solubilized with 10% SDS in 0.01N HCl. Absorbance intensities were read at 570 nm in a Tecan Infinite® 200 plate reader.

### Metabolic flux analysis and calcium imaging

BeWO cells (10,000/well) were plated into Seahorse XF96 Cell Culture Microplates (Agilent Biotechnology, Cat No. 103275-100). The oxygen consumption rate (OCR), a measure of mitochondrial respiration, and extracellular acidification rate (ECAR), a measure of glycolysis, were measured using the Seahorse XFe96 flux analyzer (Seahorse Bioscience, North Billerica, MA). At the time of assay, cell culture medium was replaced with the appropriate pre-warmed Seahorse XF Base Medium (Agilent Biotechnology, Santa Clara, CA, Cat No. 102353-100). OCR and ECAR parameters were measured using the Seahorse XFp Cell Energy Phenotype Test Kit™ (Agilent Biotechnology, Cat No. 103275-100). Metabolic stress was induced by simultaneous treatment with 1μm Oligomycin and 0.125µM Carbonyl cyanide p-[trifluoromethoxy]-phenyl-hydrazone (FCCP).

BeWO cells were also plated onto glass coverslips in 24 well plates at a density of 30,000 cells/well. After exposure to ethanol and/or forskolin in culture, cells were prepared for calcium imaging. After replacement of culture media with external imaging media (154 mM NaCl, 5 mM KCl, 2 mM CaCl_2_, 0.5 mM MgCl_2_, 5 mM glucose, 10 mM HEPES, pH 7.4), cells were loaded for 35 minutes at 37°C with the calcium indicator dye fluo-4 AM (Thermo Fisher Scientific, Cat No. F14201), at a final concentration of 5µM fluo-4 AM in 0.1% DMSO. After incubation, cells were washed to remove remaining extracellular fluo-4 and imaged at 40x using confocal microscopy (FV1200-equipped BX61WI microscope, Olympus Corporation, Center Valley, PA). Time-lapse images were acquired at a frequency of 0.5Hz. Individual cells were manually outlined and area and mean fluorescence intensity were obtained for each cell (FIJI image processing package)(105). To determine the functional calcium range of each cell, at the end of imaging, cells were exposed to 5 μM ionomycin and 10 mM EGTA (0mM external Ca^2+^, F_range_= F_ionomycin_-F_EGTA_). Baseline fluorescence was determined by averaging the lowest 5 consecutive fluorescence values during the initial 5 minutes (F_baseline_) which was then expressed as a percentage of F_range_ (ΔF_baseline_ = (F_baseline_-F_EGTA_)/F_range_ X100). Maximal intracellular calcium response to 100 µM ATP was determined by averaging the highest 3 consecutive fluorescence values during ATP application (F_ATP_) and determining the amount of fluorescence as a percentage of F_range_ (ΔF_ATP_ = (F_ATP_-F_EGTA_)/F_range_ X100).

### Quantitative reverse transcriptase-polymerase chain reaction (qRT-PCR) analysis

Total RNA was extracted from tissue, well as BeWO and HTR8 cells, using the miRNeasy Mini kit (Qiagen, Cat No. 217004). For miRNA qPCR assays, cDNA was synthesized from 200 ng of total RNA using the miRCURY LNA Universal RT cDNA synthesis kit (Exiqon, Cat No. 203301/Qiagen, Cat No. 339340, Germantown, MD) and expression was assessed using miRCURY LNA SYBR Green (Exiqon, Cat No. 203401/Qiagen, Cat No. 339345). For mRNA qPCR assays, cDNA was synthesized from 500 ng of total RNA using the qScript™ cDNA Synthesis Kit (Quanta/Qiagen, Cat No. 95047). Gene expression analysis was performed using PerfeCTa SYBR Green FastMix (Quanta, Cat No. 95073) on the ViiA 7 Real-Time PCR System (Thermo Fisher Scientific). The data presented correspond to the mean 2^-ΔΔCt^ after being normalized to the geometric mean of β-actin, Hypoxanthine-Guanine Phosphoribosyltransferase 1 (HPRT1), and 18s rRNA. Expression data for miRNA was normalized to the geometric mean of miR-25-3p, miR-574-3p, miR-30b-5p, miR-652-3p, and miR-15b-5p. For each primer pair, thermal stability curves were assessed for evidence of a single amplicon and the length of each amplicon was verified using agarose gel electrophoresis. A list of primers and their sequences is presented in Supplementary Table 1.

### Western immunoblotting analysis

Protein was extracted using 1X RIPA lysis buffer (Millipore Sigma, Burlington MA) supplemented with Halt protease inhibitor cocktail (Thermo Fisher Scientific). Tissue was homogenized using the Branson Sonifier 150. Protein concentration was determined using Pierce BCA protein assay kit (Thermo Fisher Scientific) and 30 μg of protein was loaded onto a 4-12% Bis-Tris (Invitrogen/Thermo Fisher Scientific, Cat No. NPO323BOX), size fractionated at 200 V for 35 minutes, and transferred to a PVDF membrane using the iBlot transfer system (Invitrogen/Thermo Fisher Scientific). Blots with protein from cultured cells were blocked with 5% nonfat dry milk in tris-buffered saline containing Tween®-20 (TTBS) for 1-hour and incubated overnight with primary antibody. The blot was then washed and incubated with an HRP-conjugated goat anti-rabbit or anti-mouse IgG (Invitrogen) at dilution 1:1000 for 1-hour, then developed using PerkinElmer Western Lightning Plus Chemi ECL (PerkinElmer; Waltham, MA) and visualized using a CCD camera (Fluorchem Q, Alpha Innotech; San Leandro, CA). Blots with protein from homogenized tissue were dried overnight, rehydrated in methanol, stained with REVERT™ Total Protein Stain and developed with the Odyssey CLx Imaging System (LI-COR, Lincoln, NE). Blots were then blocked with Odyssey® Blocking Buffer (TBS) for 1h and incubated overnight with primary antibody. The blot was then washed and incubated with IRDye® 800CW secondary antibody (LI-COR, Cat No. 925-32210). The following antibodies were used: β-Actin HRP (Santa Cruz Biotechnology, Cat No. sc-47778); Goat anti-Mouse IgG (H+L) Secondary Antibody, HRP (Thermo Fisher, Cat No. 62-6520); Goat anti-Rabbit IgG (H+L) Secondary Antibody, HRP (Thermo Fisher, Cat No. 65-6120); purified Mouse Anti-E-Cadherin (BD Biosciences, Cat No. 610181), Rabbit anti-vimentin antibody [EPR3776] (Abcam, Cat No. ab 924647). Protein levels were quantified using the densitometric analysis package in FIJI image processing software (105).

### ELISA

The 2^nd^ and 3^rd^ trimester maternal plasma samples were collected as part of a longitudinal cohort study conducted in two regions of Western Ukraine as part of the Collaborative Initiative on Fetal Alcohol Spectrum Disorders (CIFASD.org) between the years 2006 and 2011, as previously reported(8). Plasma, at a 1:1000 dilution, was subjected to hCG detection using Abcam’s intact human hCG ELISA kit (Cat no. ab100533) following the manufacturer’s protocol.

### Literature Review

We conducted a literature review for _HEa_miRNAs and their associated gestational pathology using the National Institute of Health’s Pubmed search interface. For each miRNA, the following search parameters were used:

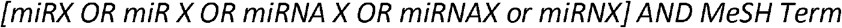

where X represents the miRNA of interest and automatic term expansion was enabled. The following MeSH terms, and related search terms (in brackets), were used: Fetal Growth Retardation [Intrauterine Growth Retardation, IUGR Intrauterine Growth Restriction, Low Birth Weight, LBW, Small For Gestational Age, SGA], Premature Birth [Preterm Birth, Preterm Birth, Preterm Infant, Premature Infant, Preterm Labor, Premature Labor], Spontaneous Abortion [Early Pregnancy Loss, Miscarriage, Abortion, Tubal Abortion, Aborted Fetus], Pre-Eclampsia [Pre Eclampsia, Preeclampsia, Pregnancy Toxemia, Gestational Hypertension, Maternal Hypertension], and Maternal Exposure [Environmental Exposure, Prenatal Exposure]. Returned articles were subsequently assessed for relevance.

### Secondary analysis of RNA sequencing data

Expression levels of _HEa_miRNAs in tissues were determined using the Human miRNA Expression Database and the miRmine Human miRNA expression database(58, 106). For expression analysis of _HEa_miRNA pri-miRNAs, RNA sequencing data was used from NCBI’s sequence read archive (https://www.ncbi.nlm.nih.gov/sra). The accession numbers for the sequence files are: uterus (SRR1957209), thyroid (SRR1957207), thymus (SRR1957206), stomach (SRR1957205), spleen (SRR1957203), small intestine (SRR1957202), skeletal muscle (SRR1957201), salivary gland (SRR1957200), placenta (SRR1957197), lung (SRR1957195), liver (SRR1957193), kidney (SRR1957192), heart (SRR1957191), whole brain (SRR1957183), adrenal gland (SRR1957124), bone marrow (ERR315396), colon (ERR315484), adipose tissue (ERR315332), and pancreas (ERR315479). Deep sequencing analysis was conducted using the Galaxy version 15.07 user interface according to the bioinformatics pipeline outlined in Supplementary Figure 1.

### Statistical analyses

Linear regression models were used to estimate associations between infant growth measures and miRNA expression levels, gestational age at blood draw, the interaction between subject-centered miRNA expression level and gestational age at blood draw, and child sex. Spearman correlations between infant growth measures and subject-centered miRNA expression levels were also calculated. Linear regression models were also used to estimate the associations between gestational at birth and log-transformed hCG levels, ethanol intake, the interaction between log-transformed hCG levels and ethanol intake, gestational at blood draw, and child sex. Statistical Analysis and graphs were generated with GraphPad Prism 6 software (GraphPad Software, Inc., La Jolla, CA), SPSS v24, or R version 3.3.1. Results are expressed as the mean ± SEM, or alternatively as box-and-whisker plots with the bounds of the box demarcating limits of 1st and 3rd quartile, a median line in the center of the box, and whiskers representing the total range of data. The overall group effect was analyzed for significance using 1-way MANOVA, 1-way or 2-way ANOVA with Tukey’s Honest Significance Difference (HSD) or Dunnett’s Multiple Comparisons post-hoc testing when appropriate (i.e. following a significant group effect in 1-way ANOVA or given a significant interaction effect between experimental conditions in 2-way ANOVA), to correct for a family-wise error rate. A 2-tailed Student’s t-test was used for planned comparisons. For experiments characterizing the individual effects of _HEa_miRNAs against the control miRNA or antagomirs, individual 2-tailed Student’s t-test with 5% FDR correction was applied to account for multiple comparisons. All statistical tests, sample-sizes, and post-hoc analysis are appropriately reported in the results section. A value of p < 0.05 was considered statistically significant, and a value of 0.1 < p < 0.05 was considered marginally significant.

### Study approval

Human study protocols were approved by Institutional Review Boards at the Lviv National Medical University, Ukraine, and the University of California San Diego as well as Texas A&M University in the USA. Research was conducted according to the principles expressed in the Declaration of Helsinki with written informed consent received from participants prior to inclusion in the study. All rodent experiments were performed in accordance with protocols approved by the University of New Mexico Institutional Animal Care and Use Committee (IACUC), and the Texas A&M University IACUC. All procedures involving non-human primate research subjects were approved by the IACUC of the Oregon National Primate Research Center (ONPRC), and guidelines for humane animal care were followed. The ONPRC abides by the Animal Welfare Act and Regulations enforced by the US Department of Agriculture.

## Acknowledgements

This research was supported by grants from the NIH, P50 AA022534 (AMA), U01 AA014835 and the Office of Dietary Supplements (CDC), R24 AA019431 (KAG), R01 AA021981 (CDK), R01 AA024659 (RCM), F31 AA026505 (AMT) and support from National Health and Medical Research Council of Australia (KMM). We thank CIFASD for intellectual support and Megan S. Pope and Tenley E. Lehman for their assistance in conducting cell culture and animal studies. Data on human subjects is deposited at CIFASD.org, in accordance with NIH data repository guidelines.

## Author contributions

AT, RM, and CC conceived of and planned the study. AT, AM, and NS designed and conducted cell culture studies, and AT conducted *in vivo*, murine miRNA overexpression studies and analyzed tissues from mouse, rat and primate PAE models. AA developed the mouse PAE model and LA and KM developed the rat PAE model and provided tissues. VR, NN, CK and KG developed the non-human primate model of PAE and provided RNA from microdissected tissues. AW and CC performed statistical analyses of human studies. AT, AM and RCM collaborated on all other statistical analyses. AT, AM, AW, NS, AA, VR, NN, CK, KG, CC, and RM collaborated in preparing the manuscript.

## Supplementary Figure Legends

**Supplementary Figure 1: Bioinformatics pipeline used to analyze _HEa_miRNA pri-miRNA expression in tissues.**

**Supplementary Figure 2: PAE paradigms in mouse, rat, and macaques**

**A)** Timeline of mouse alcohol administration

**B)** Timeline of rat alcohol administration

**C)** Timeline of macaque alcohol administration

**Supplementary Figure 3: PAE does not impair EMT in mouse placenta junctional and decidual zones**

**A)** Densitometric quantification of E-cadherin protein levels in junctional and **B)** decidual zone of control and PAE GD14 mice.

Results are expressed as expressed as the mean ± SEM, n=5-12 samples per group.

**Supplementary Figure 4: PAE and expression of core EMT transcripts in rat placenta**

**A)** Expression of CDH1, **B)** Snai1, **C)** VIM, **D)** Snai2, and **E)** TWIST in the placental labyrinth zone of PAE and control rats.

Results are expressed as expressed as the mean ± SEM, n=8 samples per group; ANOVA: significant main effect of PAE [#p<0.05].

**Supplementary Figure 5: Individual _HEa_miRNAs do not affect EMT pathway in BeWO cytotrophoblasts**

Heatmap for expression of core members of the EMT pathway following overexpression of individual _HEa_miRNAs or a control (ctrl) miRNA. Scale for heatmap coloration, right, depicts row-centered Z-score, n=10 samples per group.

**Supplementary Figure 6: _HEa_miRNAs subpools have different effect on the EMT pathway in BeWO cytotrophoblasts**

**A)** Venn diagram with the diamond indicating _HEa_miRNAs broadly implicated in gestational pathologies and the triangle outlining miRNAs implicated in preeclampsia and fetal growth restriction.

Expression of **B)** CDH1 **C)** VIM **D)** TWIST and **E)** SNAI1 transcripts following control (**C**), [hsa-miR-222-5p and hsa-miR-519a-3p] (GP), or [hsa-miR-885-3p, hsa-miR-518f-3p, and hsa-miR-204-5p] (PE/FGR) overexpression. Results are expressed as expressed as the mean ± SEM, n=5 samples per group; ANOVA: significant treatment effect [^##^p<0.05, ^###^p<0.001]. For post-hoc analysis, *p<0.05, **p<0.01, and ****p<0.0001 by Dunnett’s Multiple Comparisons.

**Supplementary Figure 7: Ethanol does not directly affect extravillous trophoblast invasion** Transwell Invasion of HTR8 Extravillous Trophoblasts following 0, 60, 120, and 320mg/dL ethanol exposure. O.D. = optical density, results are expressed as expressed as the mean ± SEM, n=8 samples per group.

**Supplementary Figure 8: Ethanol and miRNAs interfere with trophoblast cell cycle dynamics A)** Heatmap for degree of EdU incorporation in BeWO cytotrophoblasts following individual _HEa_ miRNA overexpression (top, OE) or transfection with individual miRNA hairpin inhibitors (bottom, HI). Scale for heatmap coloration, right, denotes fold change of EdU incorporation intensity relative to control mimic or hairpin transfection. N=6 samples per group, white asterisks denote _HEa_ miRNA mimics or hairpin inhibitors that had a significant effect, p<0.05,

Student’s T-test, on degree of EdU incorporation.

**B)** Degree of EdU incorporation in BeWO cytotrophoblasts following 0, 60, 120, and 320mg/dL ethanol exposure. n=5 samples per group.

**C)** Proportion of BeWO cytotrophoblasts in G_0_ /G_1_, S, or G_2_ /M phase of the cell cycle following 0, 60, 120, and 320mg/dL ethanol exposure.

Results are expressed as expressed as the mean ± SEM, n=5 samples per group; ANOVA: significant main effect of 320mg/dL ethanol exposure 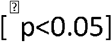. For post-hoc analysis, *p<0.05 and **p<0.01 by Tukey’s HSD.

**Supplementary Figure 9: _HEa_miRNAs influence lytic and apoptotic cell death**

**A)** Quantification of lytic cell death in BeWO cytotrophoblasts following 0, 60, 120, and 320mg/dL ethanol exposure (n=10 samples per group).

**B)** Quantification of apoptotic cell death in BeWO cytotrophoblasts following 0, 60, 120, and 320mg/dL ethanol exposure (n=8 samples per group).

**C)** Heatmap of lytic cell death in BeWO cytotrophoblasts following individual _HEa_miRNA overexpression (top, OE) or transfection with individual _HEa_miRNA hairpin inhibitors (bottom, HI). Scale for heatmap coloration, bottom, denotes fold change of lytic cell death relative to control mimic or hairpin transfection. N=10 samples per group, white asterisks denote _HEa_miRNA mimics or hairpin inhibitors that had a significant effect, <0.05, Student’s t-test, on lytic cell death.

**D)** Heatmap of apoptotic cell death in BeWO cytotrophoblasts following individual _HEa_miRNA overexpression (top, OE) or transfection with individual _HEa_miRNA hairpin inhibitors (bottom, HI). Scale for heatmap coloration, bottom, denotes fold change of apoptotic cell death relative to control mimic or hairpin transfection. N=10 samples per group, white asterisks denote _HEa_miRNA mimics or hairpin inhibitors that had a significant effect, p<0.05, Student’s t-test, on apoptosis.

**E)** Quantification of lytic cell death in BeWO cytotrophoblasts following _HEa_miRNAs or control miRNA overexpression with or without concomitant 320mg/dL ethanol exposure (n=10 samples per group).

**F)** Quantification of lytic cell death in BeWO cytotrophoblasts following transfection with _HEa_miRNA or control hairpin inhibitors with or without concomitant 320mg/dL ethanol exposure (n=10 samples per group).

**G)** Quantification of apoptotic cell death in BeWO cytotrophoblasts following miRNA mimics _Hea_ or control miRNA overexpression with or without concomitant 320mg/dL ethanol exposure (n=10 samples per group).

**H)** Quantification of apoptotic cell death in BeWO cytotrophoblasts following transfection with _HEa_ miRNA or control hairpin inhibitors with or without concomitant 320mg/dL ethanol exposure (n=10 samples per group).

Results are expressed as expressed as the mean ± SEM; ANOVA: significant main effect of 320mg/dL ethanol exposure 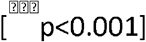, significant main effect of _HEa_ miRNA treatment [^##^ p<0.01].

**Supplementary Figure 10: _Hea_ miRNAs influence differentiation associated Ca^2+^ dynamics**

**A)** Box and whisker plot of BeWO cytotrophoblast size (left) following _HEa_ miRNA overexpression with or without 20μm forskolin treatment.

**B)** Trace of Intracellular Calcium Levels at baseline and following administration of the indicated compounds, as well as schematic and equations used to calculate relative fluorescence intensities (n=51-136 cells per group).

**C)** Box and whisker plot baseline intracellular calcium levels in BeWO cytotrophoblasts with control and _Hea_ miRNA overexpression with or without concomitant 320mg/dL ethanol exposure (n=69 to 154 samples per group).

**D)** Box and whisker plot of baseline intracellular calcium levels in BeWO cytotrophoblasts with control and _Hea_ miRNA overexpression with or without 20μm forskolin treatment (n=51 to 136 samples per group).

For box and whisker plots, bounds of box demarcate limits of 1^st^ and 3^rd^ quartile, line in middle is the median, and whiskers represent the range of data; ANOVA: significant interaction effect (sex by PAE, [^†^ p<0.05, ^† † † †^ p<0.0001]). For post-hoc analysis, *p<0.05 and ***p<0.001 by Tukey’s

**Supplementary Figure 11: Gestational age at third-trimester maternal blood collection across the UE, HEua, and HEa groups within our Ukrainian birth cohort**

Results are expressed as expressed as the mean ± SEM, n=22 to 23 samples per group

**Supplementary Figure 12: Biodistribution of miRNAs following systemic administration**

Expression of **A)** miR-518f-3p or **B)** miR-519a-3p in the indicated fetal and maternal compartments at GD12 following tail vein injection of control (NC) and miR-518f-3p or miR-519a-3p mimics (P) to pregnant C57/Bl6 dams on GD10.

n=1 sample per group, n.d. indicates non-detectable levels of miRNA.

**Table and Supplementary Table Legends**

**Table 1: _HEa_miRNAs are significantly correlated with independent measures of infant size** The correlation of 2nd and 3rd trimester maternal plasma _HEa_miRNA levels with independent measures of infant size. _HEa_miRNAs and their significantly correlated sex and gestational age-adjusted growth parameters appear in bold. *p<0.05, **p<0.01.

**Supplementary Table 1: List of primer sequences used**

**Supplementary Table 2: _HEa_ miRNAs collectively explain the variance in independent measures of infant size**

R^2^ values resulting from a multivariate statistical regression model for 2^nd^ and 3^rd^ trimester _HEa_ miRNA levels fit onto sex and age adjusted growth parameters.

**Supplementary Table 3: Maternal alcohol consumption and hCG levels are negatively correlated with gestational age at delivery**

Linear regression of gestational age at blood draw, third trimester maternal hCG levels (hCG level), degree of maternal alcohol consumption, and interaction between hCG levels and maternal alcohol consumption, with gestational age at delivery as the outcome. For maternal alcohol consumption: AAD0 and AADD0 represent absolute ounces of alcohol and absolute ounces of alcohol per drinking day around conception respectively, whereas AADXP and AADDXP represent these measures of alcohol consumption during the first trimester. Estimate represents the computed slope for each variable and C.I. is the confidence interval. *p<0.05, ***p<0.001

