## Supplementary Figures for "Maternal Circulating MiRNAs That Predict Infant FASD Outcomes Influence Placental Maturation"

### Slide 1
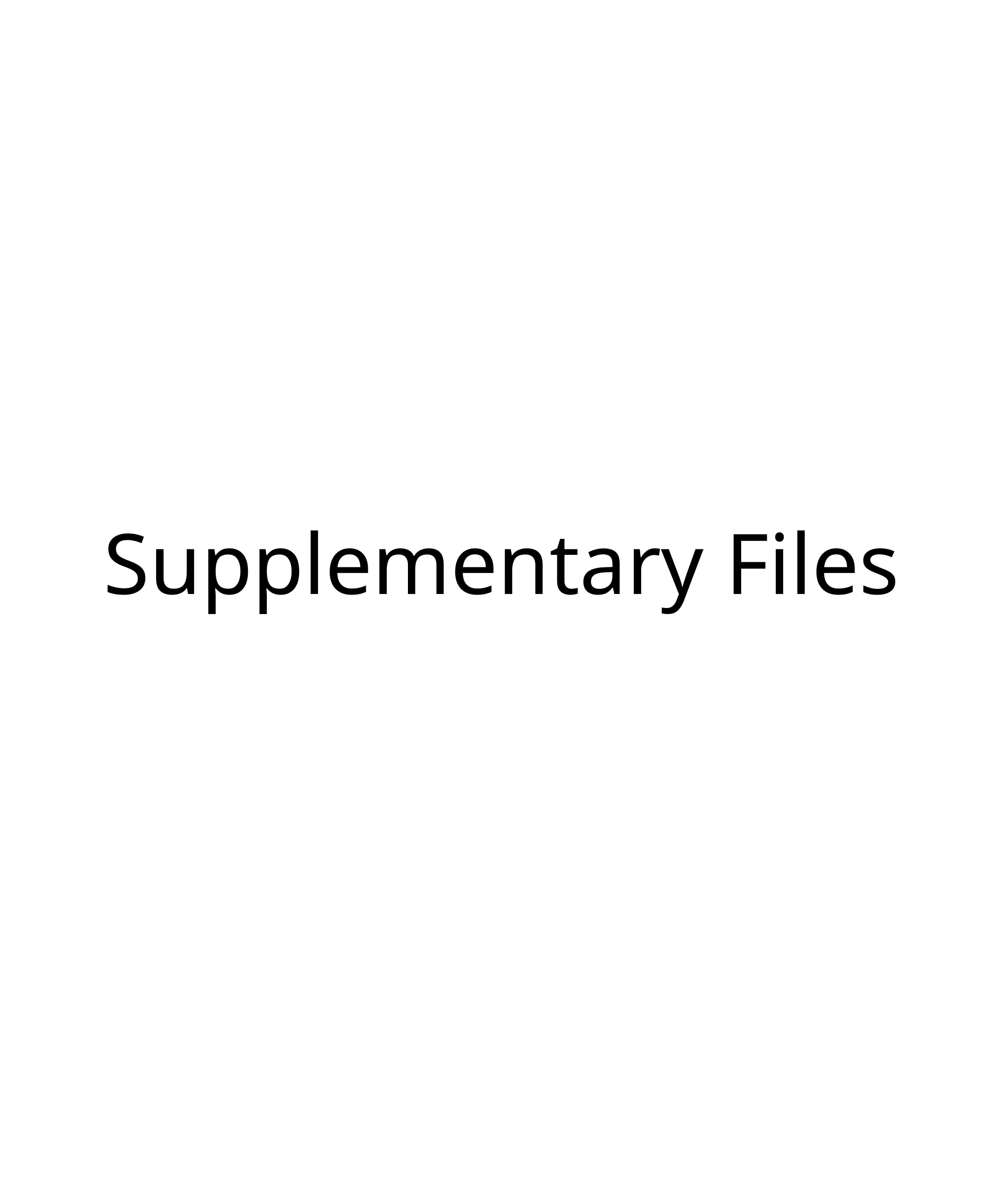

#### Supplementary Files

### Slide 2
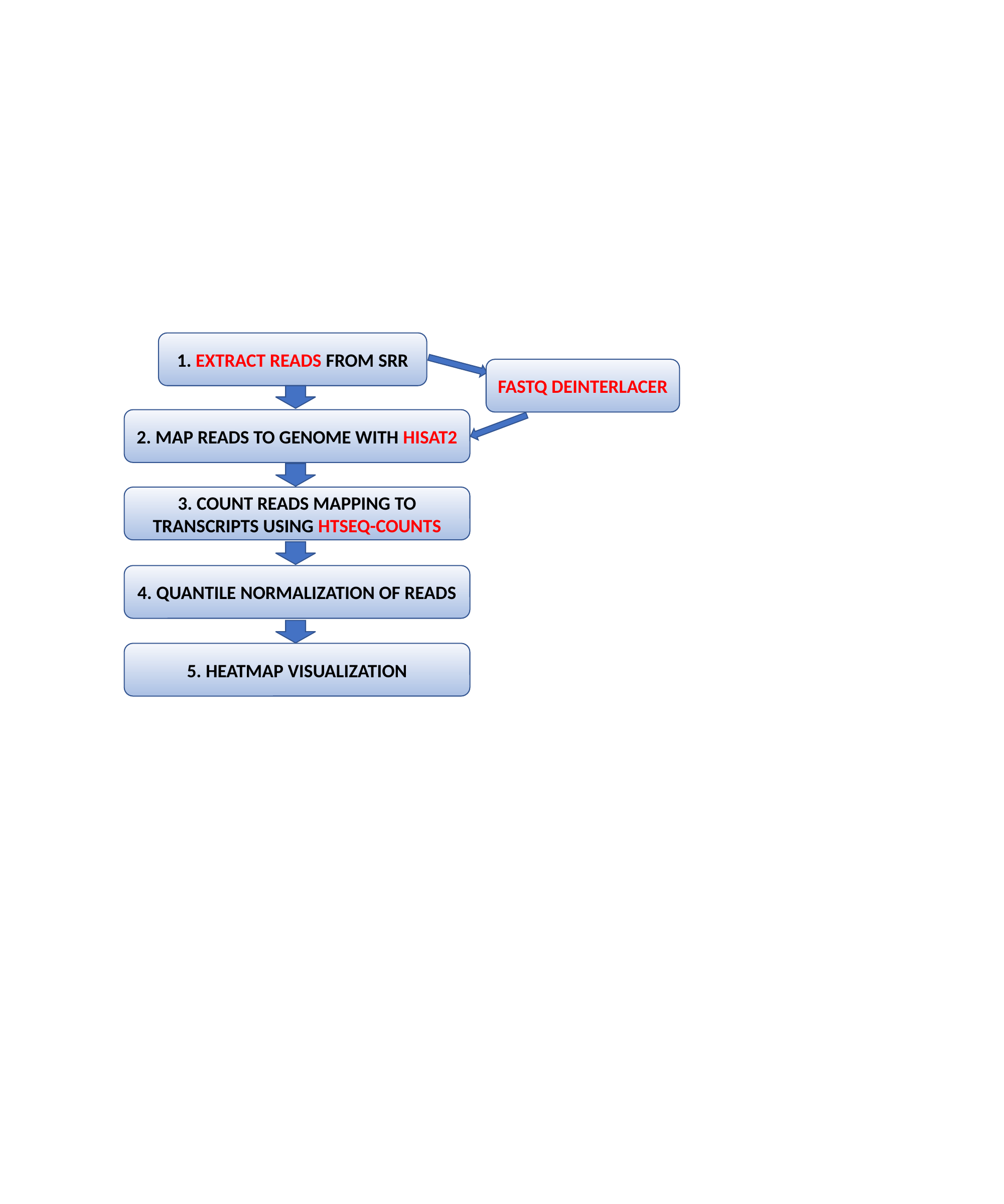

1. EXTRACT READS FROM SRR
FASTQ DEINTERLACER
2. MAP READS TO GENOME WITH HISAT2
3. COUNT READS MAPPING TO TRANSCRIPTS USING HTSEQ-COUNTS
4. QUANTILE NORMALIZATION OF READS
5. HEATMAP VISUALIZATION

### Slide 3
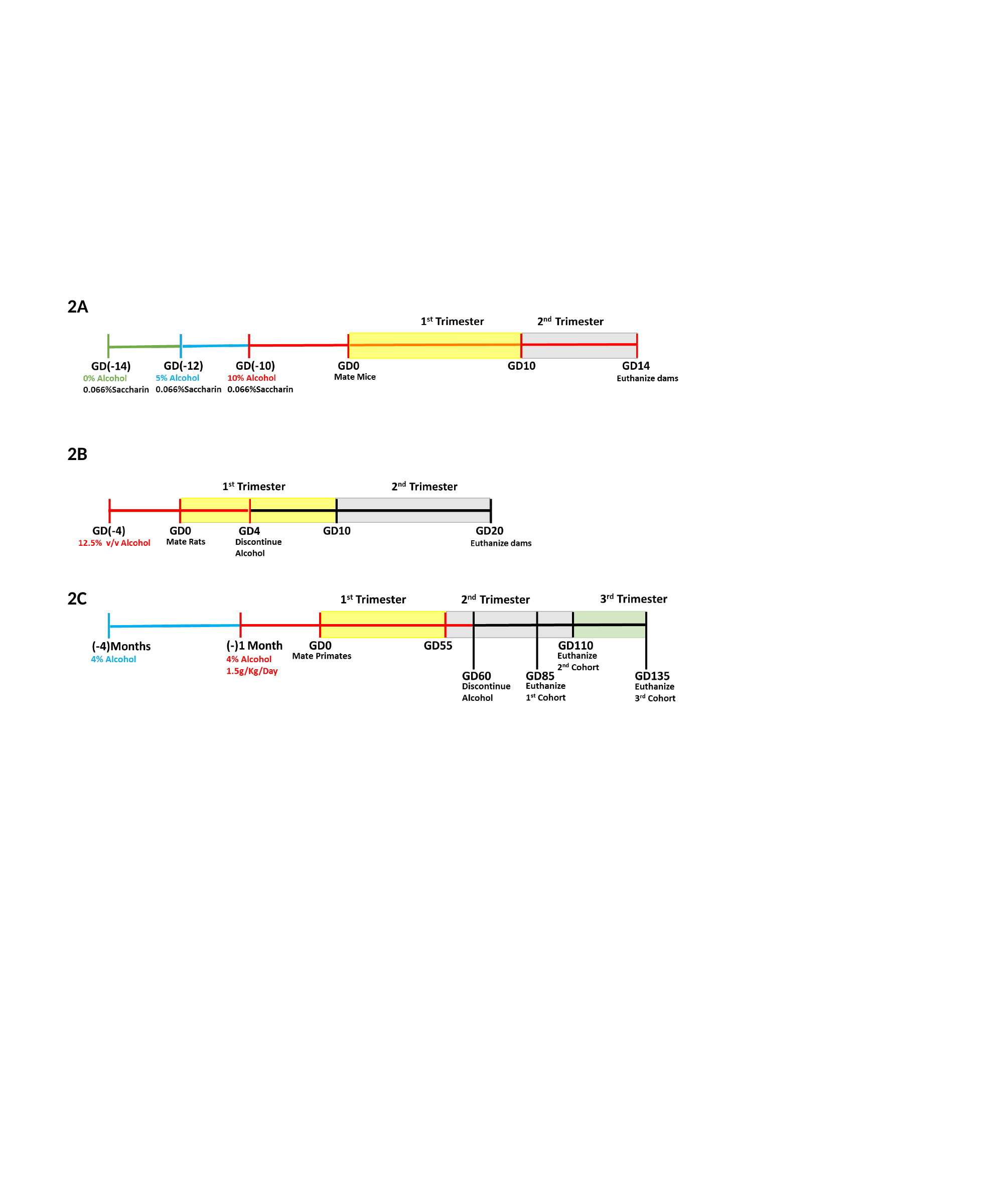

2A
2B
2C

### Slide 4
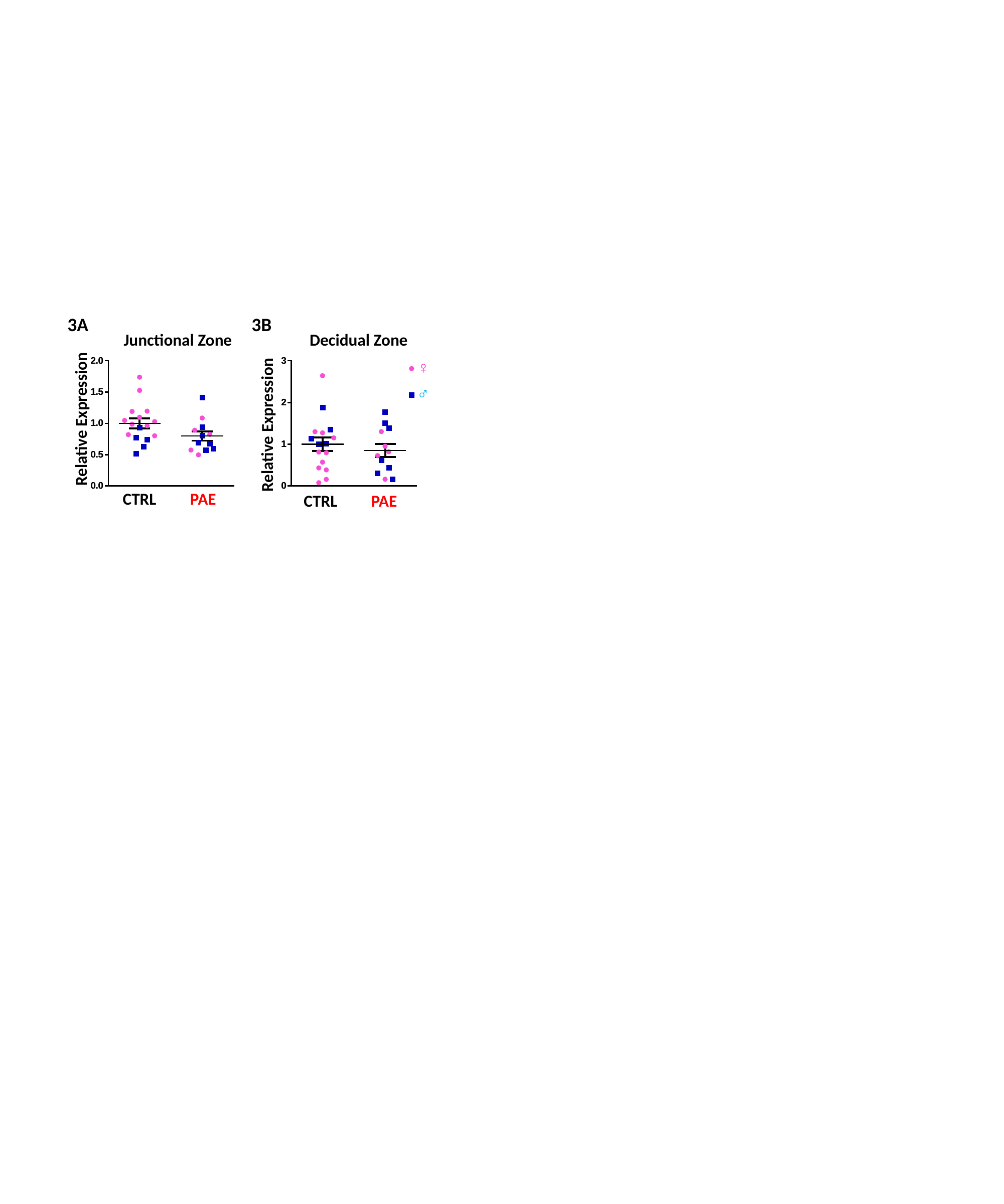

3A
3B
Junctional Zone
Decidual Zone
♀
♂
Relative Expression
Relative Expression
CTRL
PAE
CTRL
PAE

### Slide 5
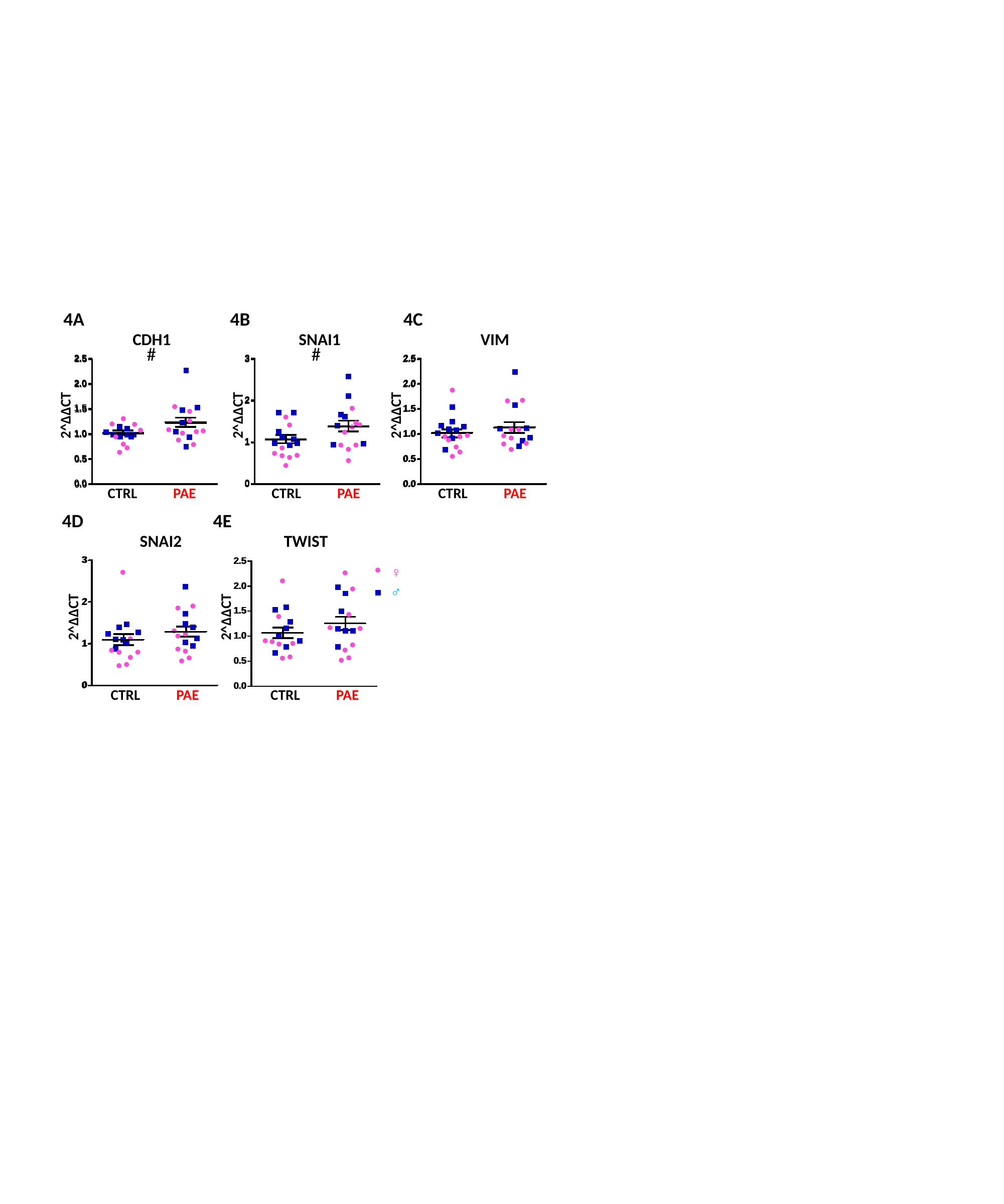

4A
4B
4C
CDH1
SNAI1
VIM
#
#
2^ΔΔCT
2^ΔΔCT
2^ΔΔCT
CTRL
PAE
CTRL
PAE
CTRL
PAE
4D
4E
SNAI2
TWIST
♀
♂
2^ΔΔCT
2^ΔΔCT
CTRL
PAE
CTRL
PAE

### Slide 6
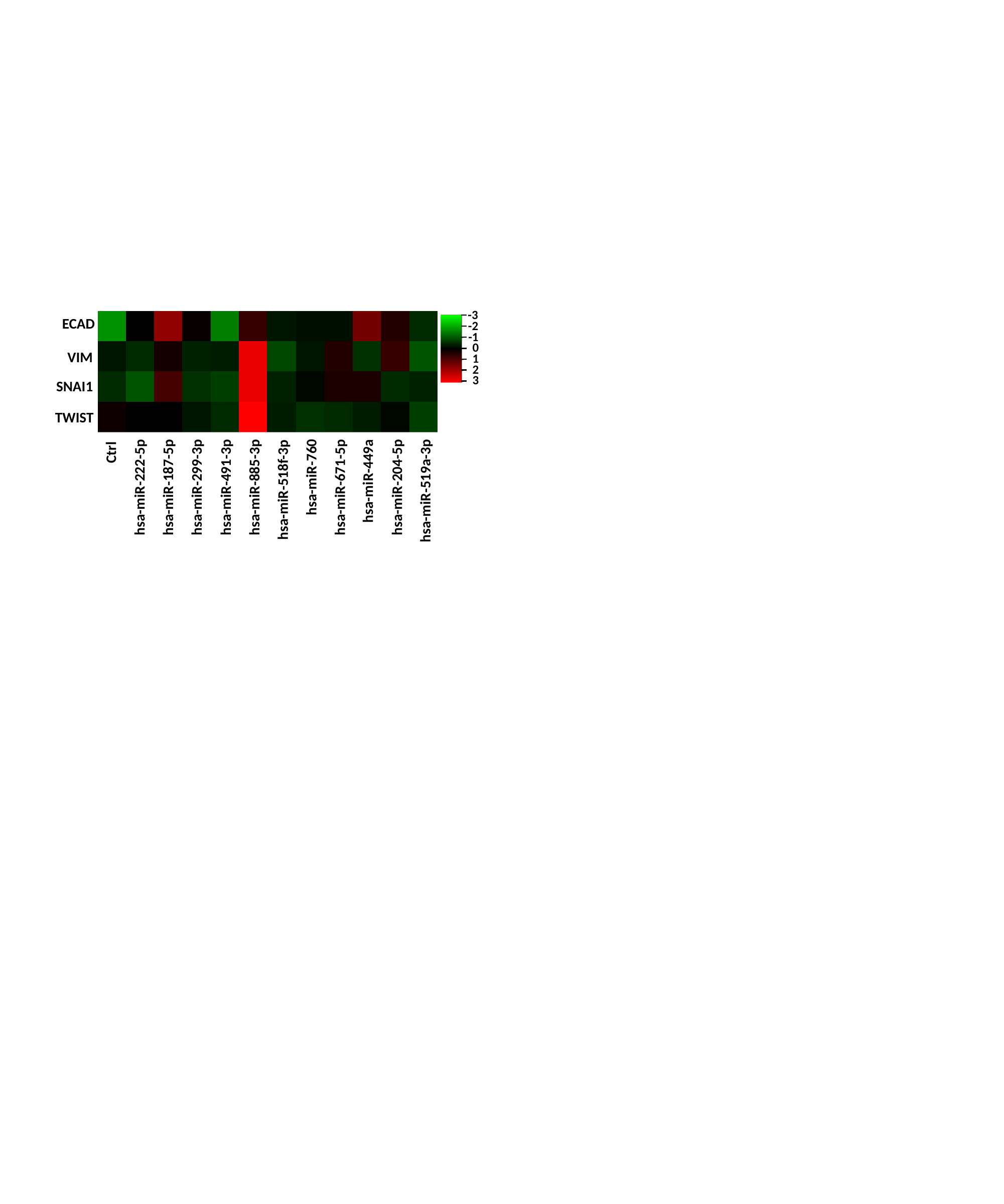

-3
ECAD
-2
-1
0
VIM
1
2
3
SNAI1
TWIST
Ctrl
hsa-miR-760
hsa-miR-449a
hsa-miR-299-3p
hsa-miR-222-5p
hsa-miR-187-5p
hsa-miR-671-5p
hsa-miR-491-3p
hsa-miR-885-3p
hsa-miR-204-5p
hsa-miR-518f-3p
hsa-miR-519a-3p

### Slide 7
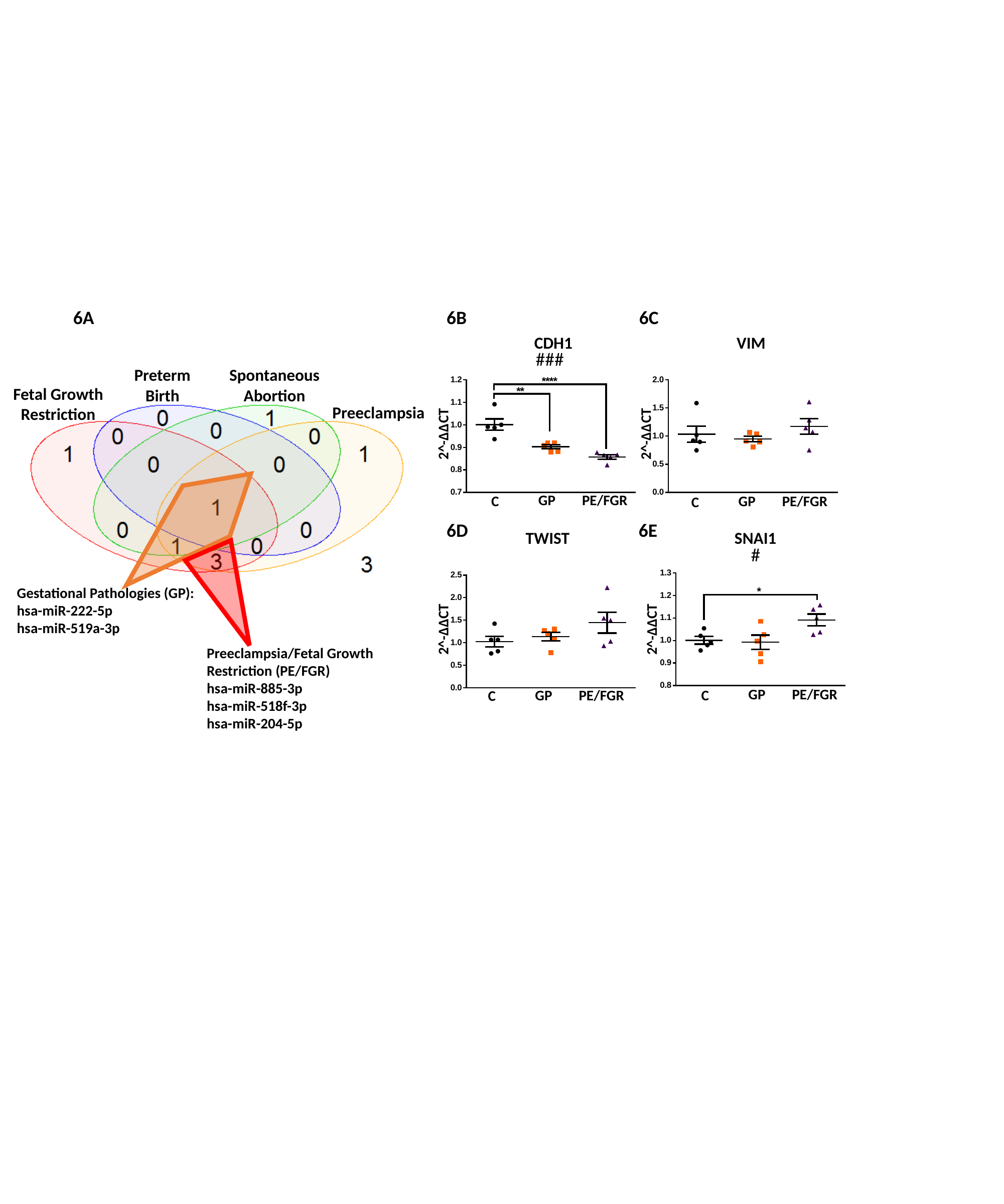

6A
6B
6C
CDH1
VIM
###
Preterm
Birth
Spontaneous Abortion
Fetal Growth Restriction
Preeclampsia
2^-ΔΔCT
2^-ΔΔCT
GP
PE/FGR
GP
PE/FGR
C
C
6D
6E
TWIST
SNAI1
#
Gestational Pathologies (GP):
hsa-miR-222-5p
hsa-miR-519a-3p
2^-ΔΔCT
2^-ΔΔCT
Preeclampsia/Fetal Growth Restriction (PE/FGR)
hsa-miR-885-3p
hsa-miR-518f-3p
hsa-miR-204-5p
GP
PE/FGR
GP
PE/FGR
C
C

### Slide 8
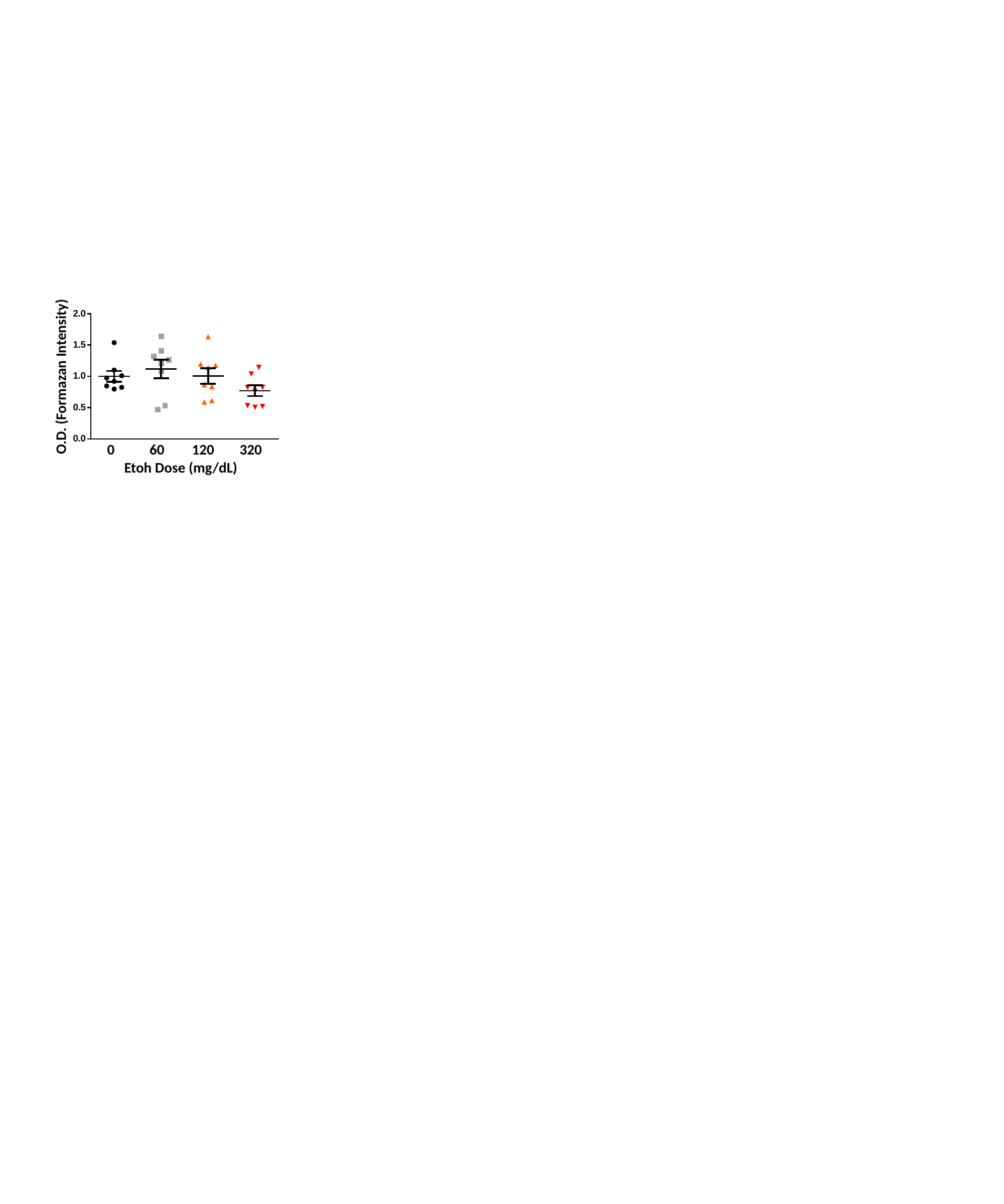

O.D. (Formazan Intensity)
0
60
120
320
Etoh Dose (mg/dL)

### Slide 9
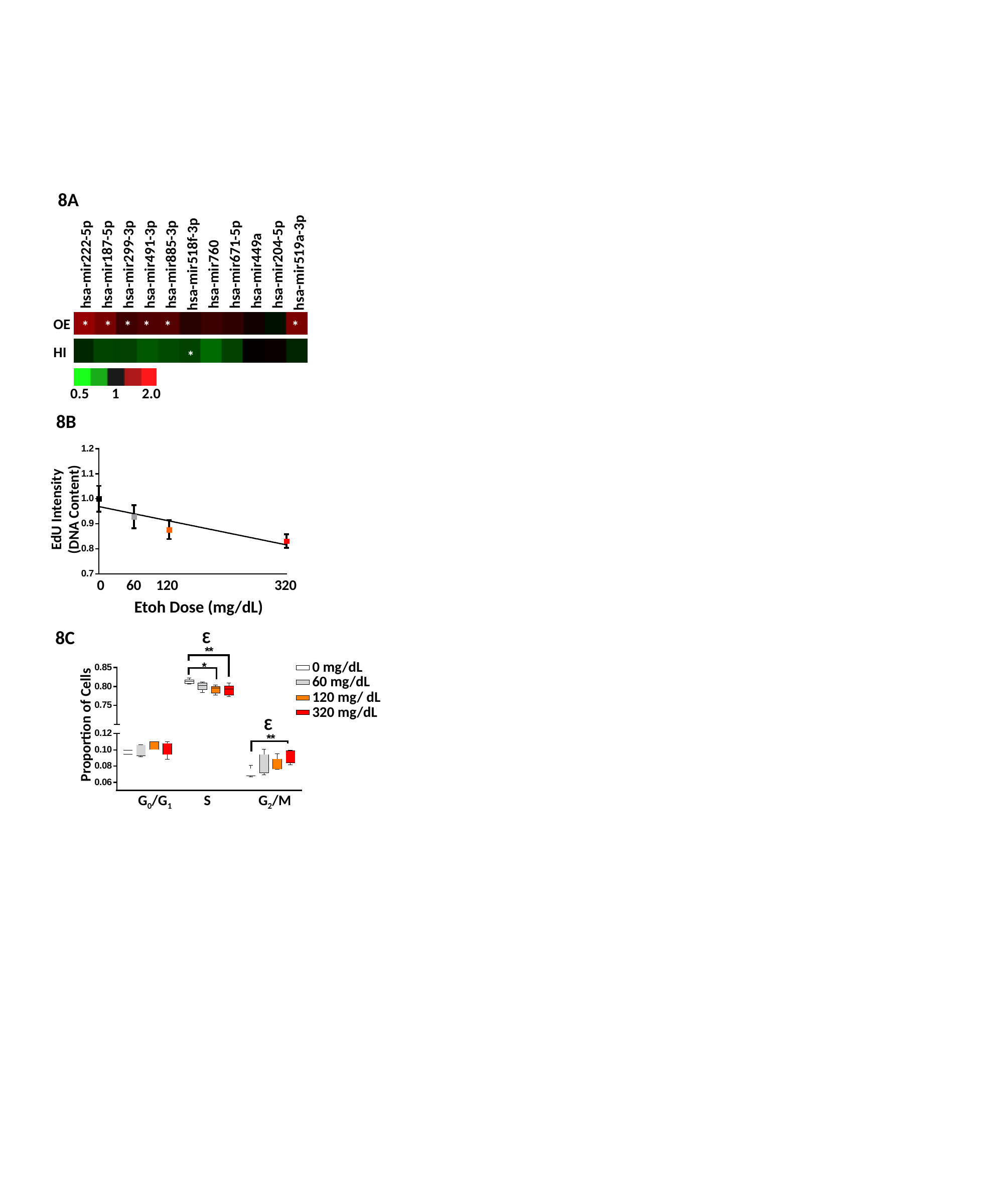

8A
hsa-mir518f-3p
hsa-mir519a-3p
hsa-mir671-5p
hsa-mir449a
hsa-mir204-5p
hsa-mir222-5p
hsa-mir187-5p
hsa-mir299-3p
hsa-mir491-3p
hsa-mir885-3p
hsa-mir760
OE
*
*
*
*
*
*
HI
*
0.5 1 2.0
8B
EdU Intensity
(DNA Content)
0
60
120
320
Etoh Dose (mg/dL)
8C
Ɛ
0 mg/dL
60 mg/dL
120 mg/ dL
Proportion of Cells
320 mg/dL
Ɛ
G_0_/G_1_
G_2_/M
S

### Slide 10
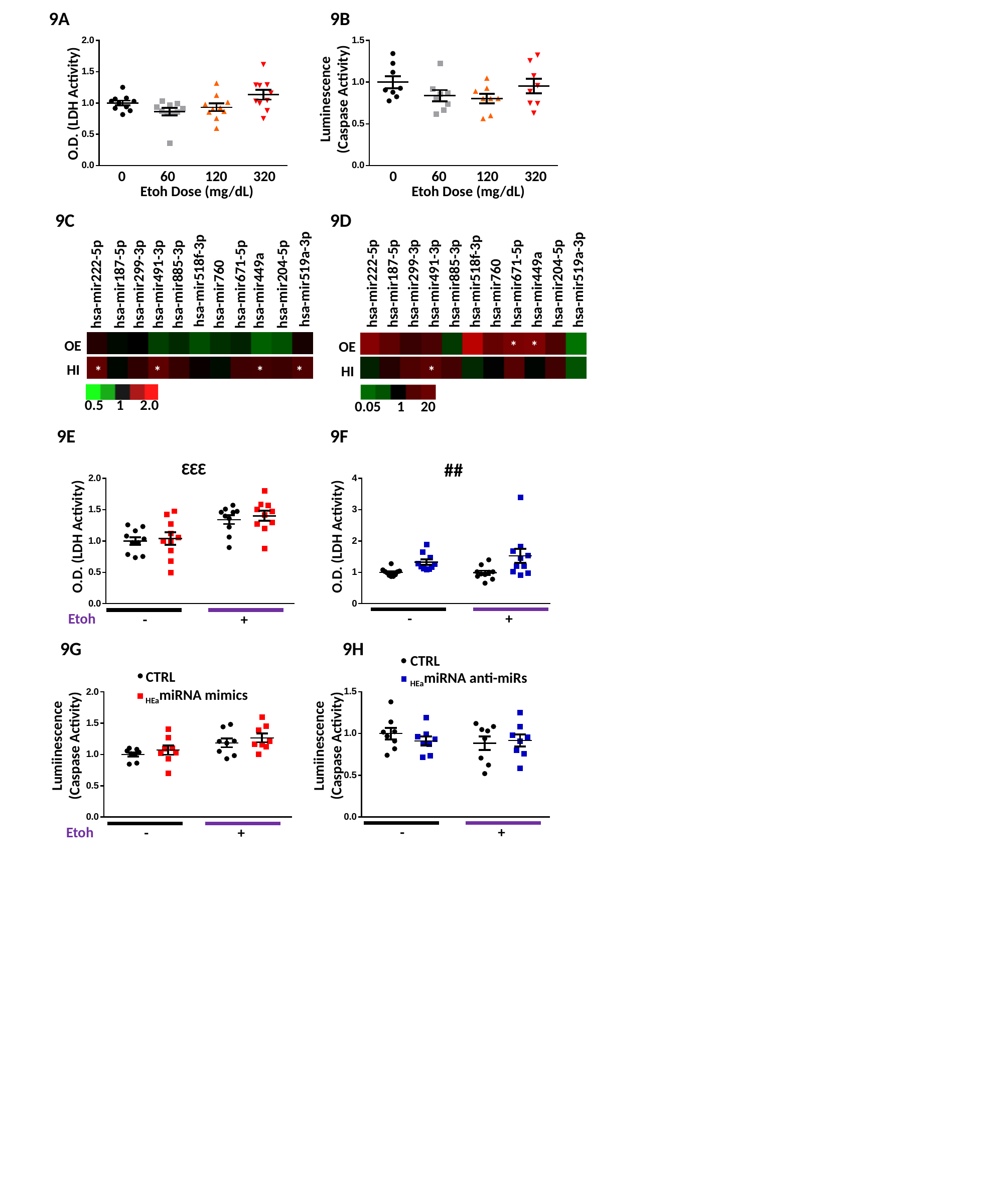

9A
9B
O.D. (LDH Activity)
Luminescence
(Caspase Activity)
0
60
120
320
0
60
120
320
Etoh Dose (mg/dL)
Etoh Dose (mg/dL)
9C
9D
hsa-mir518f-3p
hsa-mir519a-3p
hsa-mir518f-3p
hsa-mir519a-3p
hsa-mir299-3p
hsa-mir491-3p
hsa-mir885-3p
hsa-mir760
hsa-mir671-5p
hsa-mir449a
hsa-mir204-5p
hsa-mir222-5p
hsa-mir187-5p
hsa-mir299-3p
hsa-mir491-3p
hsa-mir885-3p
hsa-mir760
hsa-mir671-5p
hsa-mir449a
hsa-mir204-5p
hsa-mir222-5p
hsa-mir187-5p
OE
*
*
OE
HI
HI
*
*
*
*
*
0.5 1 2.0
 0.05 1 20
9E
9F
ƐƐƐ
##
O.D. (LDH Activity)
O.D. (LDH Activity)
-
+
Etoh
-
+
9G
9H
CTRL
CTRL
_HEa_miRNA anti-miRs
_HEa_miRNA mimics
Lumiinescence
(Caspase Activity)
Lumiinescence
(Caspase Activity)
-
+
Etoh
-
+

### Slide 11
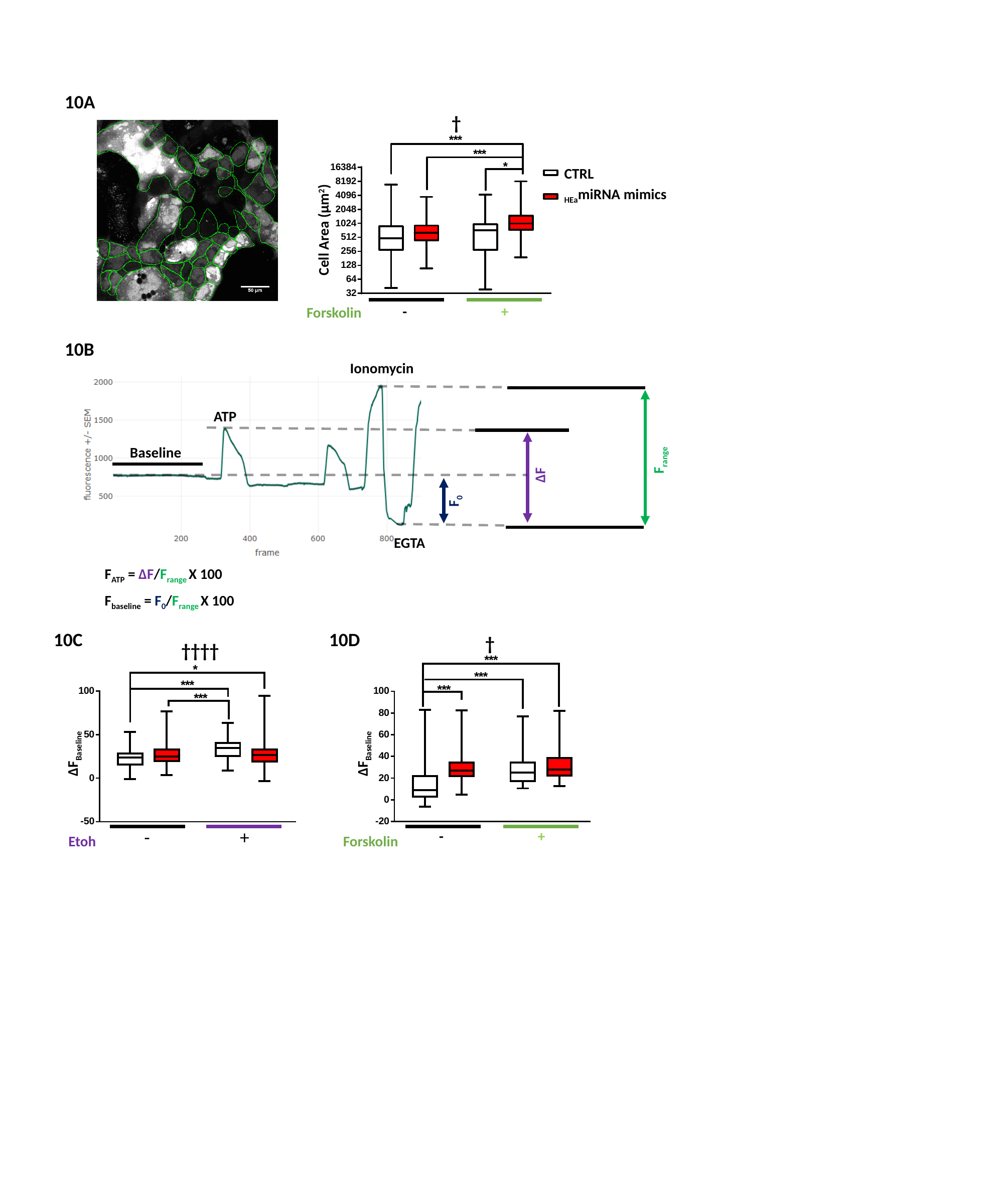

10A
†
CTRL
_HEa_miRNA mimics
Cell Area (µm^2^)
-
+
Forskolin
10B
Ionomycin
ATP
Baseline
F_range_
ΔF
F_0_
EGTA
F_ATP_ = ΔF/F_range_ X 100
F_baseline_ = F_0_/F_range_ X 100
10C
10D
†
††††
ΔF_Baseline_
ΔF_Baseline_
-
+
-
+
Etoh
Forskolin

### Slide 12
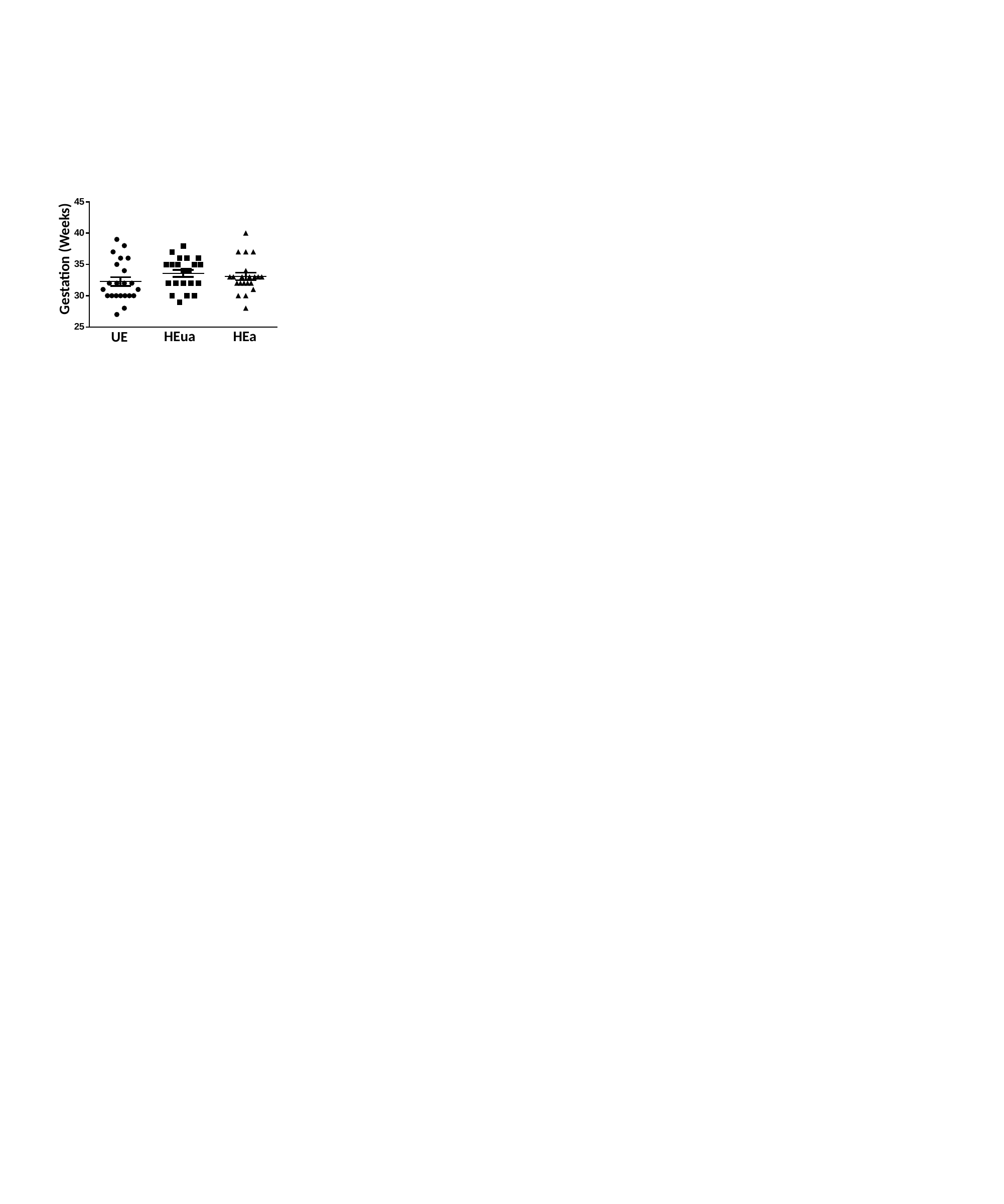

Gestation (Weeks)
HEa
HEua
UE

### Slide 13
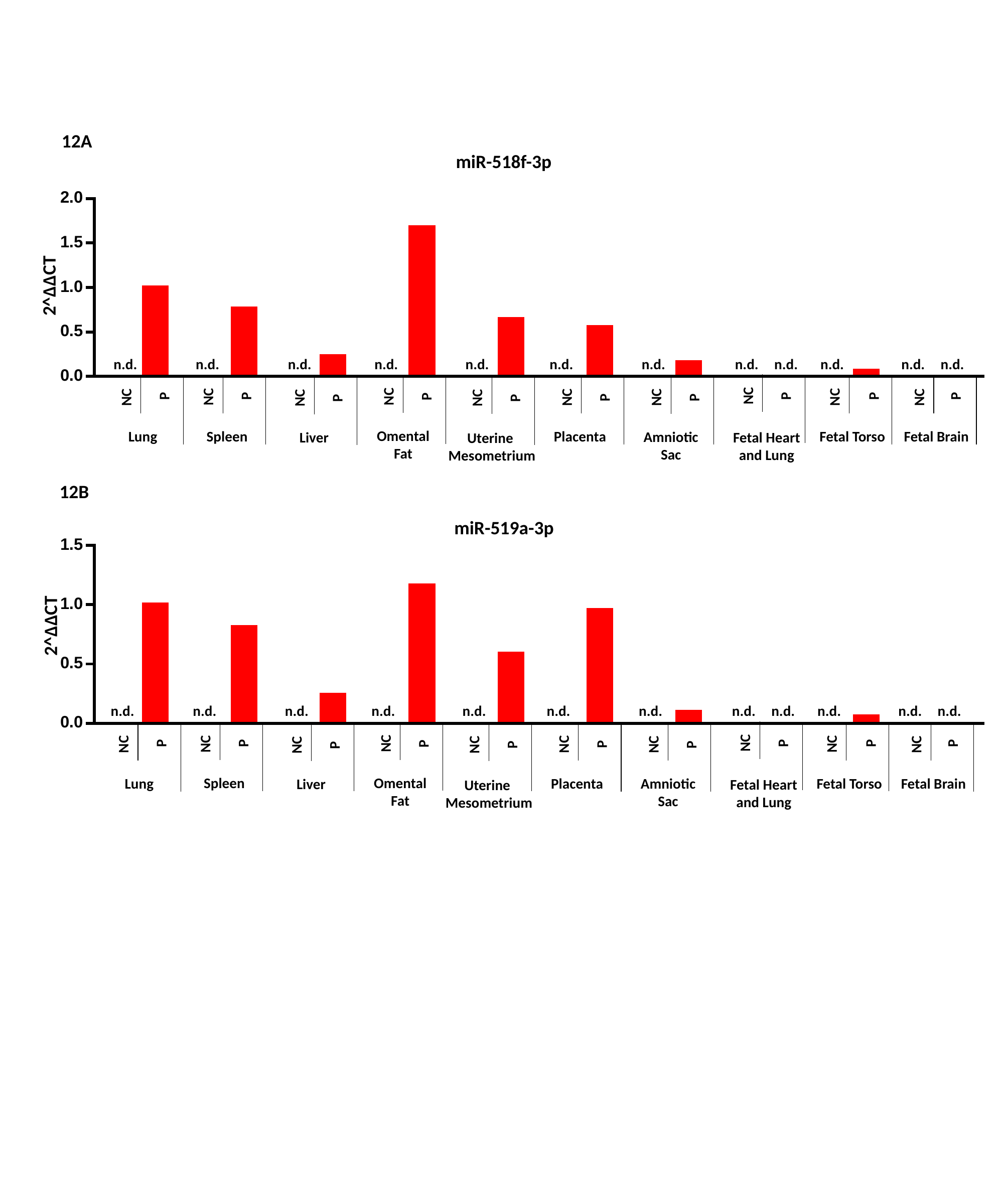

12A
miR-518f-3p
2^ΔΔCT
n.d.
n.d.
n.d.
n.d.
n.d.
n.d.
n.d.
n.d.
n.d.
n.d.
n.d.
n.d.
P
P
P
P
P
P
P
P
P
P
NC
NC
NC
NC
NC
NC
NC
NC
NC
NC
Omental
Fat
Spleen
Fetal Torso
Lung
Placenta
Fetal Brain
Amniotic
Sac
Liver
Fetal Heart and Lung
Uterine
Mesometrium
12B
miR-519a-3p
2^ΔΔCT
n.d.
n.d.
n.d.
n.d.
n.d.
n.d.
n.d.
n.d.
n.d.
n.d.
n.d.
n.d.
P
P
P
P
P
P
P
P
P
P
NC
NC
NC
NC
NC
NC
NC
NC
NC
NC
Omental
Fat
Spleen
Fetal Torso
Lung
Placenta
Fetal Brain
Amniotic
Sac
Liver
Fetal Heart and Lung
Uterine
Mesometrium
